# Deep analysis of FANTOM CAGE data reveals hierarchical patterns of TSS co-deployment hubs and their disruption in cancers

**DOI:** 10.64898/2026.05.15.725323

**Authors:** Ruthwick Meduri, Aditi Lakshmi Satish, Umashankar Singh

**Affiliations:** HoMeCell Lab, Department of Biological Sciences and Engineering, Indian Institute of Technology Gandhinagar, Gujarat, 382055 India

**Keywords:** CAGE, Transcription start site, Transcriptional covariance, CTCF, YY1, Cancer

## Abstract

Selective deployment of multiple transcription start sites is a major regulatory feature of human transcriptomes. FANTOM CAGE data exhibit a near-universal TSS deployment parsimony which is disrupted in cancers. We have recently shown that TSS deployment is sensitive to gene function, futile upstream transcription, and cellular biosynthetic states. Patterns in FANTOM CAGE data can reveal mechanisms underlying TSS co-deployments. We propose and test the possibility that some TSSs act like epromoters and act as co-varying hubs of transcriptional activities for multiple other promoters. Using deep analysis of CAGE data implemented through neural networks we show that non-cancers implement transcription co-deployments through cores of epromoter-like TSSs which are generally proximal to their start codons. These TSSs show enhancer-like TFBSs profiles. A comparison with cancer CAGE data shows that the concentrated epromoter core is disrupted in cancers with multiple distal TSSs replacing the proximal TSS cores. We provide evidence that the core TSSs are rich in YY1 and CTCF binding sites and associated with genes coding for transcription factors. Our findings show that covariance of TSS deployment is sensitive to transcriptional resource cost and a parsimonic design of TSS co-deployments depends on proximal TSSs in non-cancers, a mechanism grossly disrupted in cancers.

**Highlights:**

- Heterogeneous FANTOM CAGE data contains universal patterns of TSSs co-deployments.
- TSS co-deployments exhibit a parsimonious “core-covariant” scheme which is disrupted in cancers.
- Core TSSs are enriched in transcription factor binding sites and gene functions which justify biological features of the samples.
- The DL pipeline we present identifies the core-covariant TSS sets in an unbiased manner.

## 1 Introduction

Alternative TSS deployment is a major mechanism determining transcriptomic diversity and complexity in metazoans with consequences on cellular functioning and fitness (Zhan et al. 2025; Carninci et al. 2006; Preker et al. 2008; Core, Waterfall, and Lis 2008; Kimura et al. 2006). TSS selection is tissue type specific (Moore et al. 2022; Zhou et al. 2022; Hou, Hon, and Huang 2023). Genes have dominant or major promoters and start sites and the selection of dominant start sites can vary between cell types (Carlos Alfonso-Gonzalez et al. 2023; Li et al. 2019; Xu, Park, and Zhang 2019; Zhan et al. 2025; Yu et al. 2017; Duttke et al. 2024). Coordinated transcription of genes, executed through coordinated TSS deployments, is essential for cellular identity and functioning. Although TSSs are structurally autonomous units with DNA sequence properties under mutually independent mutational and evolutionary pressures, their deployments are co-regulated (Karlsson, Lönnerberg, and Linnarsson 2017; Barozzi et al. 2014; Wiechens et al. 2025). TSS deployments are sensitive to the availability of resources such as transcription factors, RNA polymerases, associated co-factors and nucleotides (Kopytek and Peterson 1998; Conaway and Conaway 1988; Dvir et al. 1996; Rappaport and Weinmann 1987; Sawadogo and Roeder 1984; Noviello 2025; Traut 1994). In addition, TSS deployment is regulated by mechanisms including transcription factor binding sites in proximity of the TSSs, local epigenotypes such as histone modifications and cytosine methylation patterns as well as spatiotemporal chromatin topologies (Barbadilla-Martínez et al. 2026; “Evolution of Diverse Strategies for Promoter Regulation” 2021).

TSS have been broadly classified as sharp tissue type specific TATA-box containing TSSs or TATA-less CpG-rich broad TSSs with near-ubiquitous activity (Serebreni et al. 2023; Valen and Sandelin 2011; Ponjavic et al. 2006). The major/minor TSS identities are subject to the tissue type being analyzed. TSS switches are fundamental to zygotic gene activation (Schulz and Harrison 2019) and deregulated in cancers (Demircioğlu et al. 2019; Zhao et al. 2024; Bone and Inman 2025; Thorsen et al. 2011). It follows that some locally acting genomic features differ between TSSs and determine their background deployment potentials as major or minor TSSs (C. Alfonso-Gonzalez and Hilgers 2024; Fu and Li 2025). Predictably, neighboring and nested TSSs are likely to be similarly affected by such local elements and have similar background deployment potential (Carninci et al. 2006; Juven-Gershon and Kadonaga 2010). This is further supported by the observations that neighboring genes have correlated expression patterns assuming that most TSSs of a gene are located closer to each other than those of a physically distant gene with uncorrelated expression (Ghanbarian and Hurst 2015; Michalak 2008; Purmann et al. 2007; M Ribeiro, Ziyani, and Delaneau 2022; Dong, Brown, and Truong 2024; Zinani, Keseroğlu, and Özbudak 2022). Thus, local DNA sequence-dependent deployment potential of TSSs would be primarily autonomous with the deployment prone to molecular errors and stochasticity (van Arensbergen et al. 2017; Weingarten-Gabbay et al. 2019; Dudnyk et al. 2024); this autonomous property of the TSSs would serve as an input to regulatory epigenetic and biochemical mechanisms. Enhancers can transactivate TSSs through *cis* proximity spatiotemporal colocalization or facilitation of co-activator access in active hubs of phase separated condensates (Buitrago, Vidal, and Lim 2026).

Correlated activities of genes located on different chromosomes have been observed (Van Dyke et al. 2021; Zaborowski and Walther 2020; Lan et al. 2006; Albert et al. 2018). However, the coordinated TSS deployment of TSSs in *trans* is sensitive to the influence of DNA sequences; noteworthily, eQTLs affect gene expression coordinatedly in *trans* (Yao et al. 2017). Correlated TSS deployment can also result from common DNA-binding proteins, such as CTCF and YY1 (Verheul et al. 2020; Dehingia et al. 2022), which bring distant TSSs into a complex through less rigid structure and interactions (Tan and Ma 2025; Harmston and Lenhard 2013; Phanstiel and Wang 2022; Woodworth and Lakadamyali 2024). Coordinated TSS deployments in topological clusters operate as a “one enhancer-multiple promoter” system (Uyehara and Apostolou 2023; Buitrago, Vidal, and Lim 2026; Perlman et al. 2024; Zhu et al. 2021) with some enhancers being robust promoters themselves (Zhu et al. 2021; Paramo et al. 2026; Medina-Rivera et al. 2018; Malfait, Wan, and Spicuglia 2023; Wan et al. 2025). The one-to-many hierarchy of TSSs in such a setup shall be potentially identifiable with TFBS distributions such that driver TFBSs, such as those of YY1, concentrated at the central and deterministic “one” component (Weintraub et al. 2017) whereas degenerate cooperating TFBSs present at the “many” peripheral TSSs.

CAGE data reports promoterome out of many human tissues. Satish *et al* (Satish et al. 2025) curated them into non-cancers or cancers and reported near-universal features of non-cancer promoteromes disrupted in cancers. TSSs concomitant with a preferential proximal TSS deployment for cancer hallmark-supporting genes. The TSS deployment patterns in non-cancers are not expected to generate such parsimonious patterns stochastically or autonomously. We propose and test, using a deep analysis of CAGE data implemented through neural networks, the possibility that some TSSs act like epromoters and act as hubs of transcriptional activity covariance for multiple other promoters. A recent report by Satish and coworkers (Satish et al. 2025) has defined such generic statistical features of TSS deployment in non-cancer differentiated cells which are perturbed in cancers.

TSS deployments, as represented in CAGE data, have several embedded properties which differ between cancers and non-cancers. The TSS deployment rates exhibit significantly different entropies in gene-space (Shannon entropy of TSS usage for a gene across multiple samples) and sample-space (Gene-independent Shannon entropy of TSS usage across multiple samples) leading to an overall higher futile transcription in cancers. This futile transcription, defined as the net ribonucleotide count transcribed between the TSS and the start codon, is increased in cancers by either a leaky transcription from distant upstream TSSs or high transcription rates from proximal TSSs. Interestingly, the latter TSSs with high transcription from proximal TSSs belong to genes involved in biosynthesis and chromatin organization. We propose that the “one-to-many” co-deployment model shall be predictable using a combination of the futile transcription distance, TSS deployment rates and associated entropies. The “one-to-many” co-deployment model shall be parsimonious and normally utilize TSSs with less futile transcription distance as the “one” central co-variant to multiple partner TSSs with no futile transcription constraint on the latter. The model shall be predictably different between cancers and non-cancers and justify the functional categories of genes which are dysregulated in cancers.

We test the existence of such a parsimonious “one-to-many” model in which TFBS-driven, resource sensitive and gene function responsive co-deployments of TSSs define general principles of coordinated gene expression. By utilizing CAGE data, we develop deep clustering pipelines which reveal such “one-to-many” models in differentiated non-cancer samples and its variants in cancers. By analyzing the CAGE data and genomic features of the TSSs reported by our pipeline we show that the “one-to-many” co-deployment model exists generically in non-cancers with a predominant effect of parsimonious negative covariance between pairs of TSSs. The cancers on the other hand show a variation of this model in which the parsimony is lost with specific targeting of genes supporting cancer hallmarks, specifically RNA biosynthesis.

## 2 Materials & Methods

### 2.1 Curation of CAGE TSSs and their derived parameters

For our analysis, we curated a comprehensive set of 55,048 transcription start sites (TSSs) and their associated CAGE derivatives. All parameters utilized in this study, including the curated set of TSSs, their scaled scores (γ) and the derived metrics which includes distance between transcription start site to the most upstream start codon (*d*), transcript count index (*TCI*), energy cost coefficient (*E*), base count index (*BCI*), effective *d* (*vd*), gene–space entropy (*gH*),sample–space entropy (*sH*) were generated following the methodology established by Satish and co-workers (Satish et al. 2025). These curated features serve as the inputs for the downstream pairwise covariance analysis and the training of the data-specific denoising autoencoders.

### 2.2 Pan-promoterome TSS pairing and covariance calculation

All curated TSSs were used to generate exhaustive pairs, represented as *TSS_A_•TSS_B_*. These pairs were structured such that their distances from the TSS to the most upstream start codon (*d*) satisfied the condition, *d_TSSA_*> *d_TSSB_* where *d_TSSA_* and *d_TSSB_*are *d* of *TSS_A_* and *TSS_B_* respectively. Self-pairs and *TSS_B_•TSS_A_* pairs were eliminated. This resulted in the curation of approximately 1.5×10^9^ unique *d*-limited TSS pairs for further investigation. This was achieved by making an unbiased 55,048×55,048 TSS pair matrix followed by removal of diagonal pairs and *TSS_B_•TSS_A_* pairs.

Covariance was then calculated for each TSS pair based using scaled CAGE scores. Covariance was calculated exclusively using samples where both TSSs in a given pair exhibited non-zero scaled scores; if a score of zero was present for either TSS in a specific sample, that sample was eliminated from the calculation of covariance for that pair.

### 2.3 Training and implementation of denoising autoencoder

#### 2.3.1 Feature Engineering for Model Training

Input features for the denoising autoencoder were constructed by integrating multiple CAGE-derived parameters, including scaled expression scores, *d*, and entropy. There are 15 specific features (not shown) which we have utilized for model training. These engineered features were designed to capture the complex regulatory landscape and spatial relationships inherent in the TSS datasets. Following data filtration, a random subset comprising 10% of the total TSS pairs (1.5×10^9^) was selected to constitute the final training dataset.

#### 2.3.2 Architecture of the denoising autoencoder

To reduce dimensionality and extract robust biological signals from the high-dimensional TSS pair feature space, we designed and implemented a bottleneck autoencoder architecture. The model was developed using the PyTorch framework and optimized for high-performance computing environments. The primary objective of this architecture was to compress the 15-dimensional input vector, comprising pairwise covariance, distance ratios, and CAGE-derived metrics, into a compact 4-dimensional latent representation while minimizing reconstruction error.

The autoencoder consists of a symmetrical encoder-decoder structure. The encoder transitionally reduces the dimensionality through a series of fully connected layers, beginning with a 12-node stabilization layer, followed by an 8-node intermediate layer, and culminating in a 4-node latent bottleneck layer. Each linear transformation in the encoder is followed by Batch Normalization to stabilize internal covariate shift and a Parametric Rectified Linear Unit (PReLU) activation function, allowing the model to adaptively learn the optimal slope for negative activations. The decoder mirrors this architecture, projecting the latent features back through 8-node and 12-node layers to reconstruct the original 15-dimensional input.

#### 2.3.3 Model Training and Optimization Strategy

Prior to training, the dataset was shuffled and partitioned into a 90% training set and a 10% test set for validation. To ensure uniform feature contribution, a standard scaling transformation was computed exclusively on the training data and subsequently applied to both partitions. The model was trained using the Adam optimizer with an initial learning rate of 10^-5^ and a weight decay of 5×10^-5^ to provide L2 regularization. We utilized a Mean Absolute Error (L1) loss function to supervise the reconstruction process, as it provides greater robustness to outliers within the genomic feature distribution compared to standard squared error metrics.

Training was conducted over a maximum of 100 epochs with a batch size of 524,288 samples to leverage GPU parallelization. To optimize convergence, we employed a Cosine Annealing learning rate scheduler and implemented a gradient clipping threshold of 1 to prevent exploding gradients. An early stopping mechanism with a patience of 5 epochs was utilized to monitor validation loss, ensuring that the final model parameters were selected from the state of optimal generalization before the onset of overfitting. The final latent representations extracted from the bottleneck layer were preserved for downstream clustering and biological interpretation.

To investigate transcriptional differences between physiological states, two independent denoising autoencoder models were trained using distinct subsets of CAGE data. These subsets were categorized as either “non-cancer” or “cancer” based on the classification methodology reported by Satish et al. By training separate models on these state-specific samples, we were able to capture the unique regulatory architectures and covariance patterns characteristic of both non-cancer and cancer data.

### 2.4 Extraction and clustering of latent vectors and core-TSS identification

Following the filtration and feature extraction processes described above, feature vectors for all TSS pairs from both non-cancer and cancer samples were subjected to the denoising autoencoders to extract latent representations. Each dataset non-cancer (*N*) and cancer (*C*) was processed through both the non-cancer and cancer models.

The final analysis incorporated four distinct combinations: these represent non-cancer data processed through the non-cancer model, cancer data through the non-cancer model, cancer data through the cancer model, and non-cancer data through the cancer model. This approach allowed for the systematic extraction of latent vectors across all cross-conditional configurations.

Independent and exhaustive *k*-means clustering was performed on latent vectors obtained from each configuration across a range of *k* values from 3 to 20. For each iteration, the resulting cluster compositions were evaluated based on four primary metrics: the total number of TSS pairs, the number of unique high-*d* TSSs, and the number of unique low-*d* TSSs. To determine these unique counts, the TSS pairs within each cluster were dissociated into their individual components and quantified.

The optimal number of clusters (*k*) was selected by identifying the minimum *k* value that produced the first cluster with a minimum of 100,000 TSS pairs characterized by a disproportionately low unique count of either high-*d* or low-*d* TSSs. Within these selected clusters, the population with the lower unique TSS count was defined as core TSSs, while the population with the higher unique count was designated as covariant TSSs.

To facilitate a comparative analysis of state-specific transcriptional regulation, we identified exclusive TSS populations unique to each model•data configuration. These exclusive sets were defined by isolating TSS pairs that appeared uniquely within one specific pairing of a trained model (Normal or Cancer) and its input dataset (Normal or Cancer). For instance, N•N and N•C represent TSS pairs exclusive to normal and cancer data, respectively, when processed through the normal model; similarly, C•N and C•C denote the exclusives derived from the cancer model. This model•data based nomenclature is followed all throughout.

### 2.5 Visualization of Core Covariance Interactions

The core covariance interactions were visualized using Circos (v0.69-8) (Krzywinski et al. 2009) to represent the genomic distribution of the identified pairs. In these plots, links were generated to connect the core TSSs with their corresponding covariant TSSs. To facilitate the identification of inter- and intra-chromosomal interactions, the color of each link was assigned based on the specific color of the chromosome to which the covariant TSS belongs.

### 2.6 Rank-based analyses of TSSs

#### 2.6.1 Covariant count-rank

To quantify the influence of each core TSS within the core-covariant networks, we ranked them based on their interactions, defined by the total number of associated covariant TSS partners.

Core TSSs were ranked in descending order, such that the TSS with the highest number of covariant partners received the top rank, while the TSS with the fewest partners was ranked last. In instances where multiple core TSSs possessed an identical number of covariant partners, they were assigned the same rank.

These rankings were utilized in two distinct contexts. The ranks were represented as standalone values without further modification. Conversely, when used as coordinates on the X-axis the ranks were partitioned into percentiles to facilitate analysis. In this case, the corresponding Y-axis values falling within each percentile was aggregated by calculating their total sum.

#### 2.6.2 Core d-rank

To represent the relationship between core TSS *d* and associated functional parameters, we implemented a percentile-based ranking and aggregation strategy. Initially, all unique core TSSs were ranked according to their *d* in descending order, such that the TSS with the highest *d* value received the top rank. Following this primary ranking, the corresponding Y-axis parameters were aligned with their respective *d*-ranks. In cases where multiple Y-axis values shared a specific *d*-rank, these values were either summed or averaged to produce a single representative value for that rank.

To facilitate visualization, the *d*-ranks were subsequently partitioned into percentiles. The Y-axis values falling within each specific percentile were then aggregated, either by calculating the total sum or the mean, to generate the final data points for the respective figures.

#### 2.6.3 Covariant sum-rank

To analyze the relationship between core TSS *d* and the total connectivity of the core-covariant network, we implemented a combined ranking and summation approach. As previously described, core TSSs were first ranked by their *d*, and these ranks were partitioned into percentiles to form the X-axis. For each *d*-percentile, we calculated the aggregate association by summing the total number of covariant TSS partners associated with all core TSSs falling within that specific percentile.

These cumulative sums were then subjected to a secondary ranking in descending order, where the percentile containing the highest aggregate number of covariants received the top rank and the percentile with the lowest sum received the bottom rank. This resulting ranked parameter was plotted on the Y-axis against the core-*d* percentile ranks on the X-axis.

#### 2.6.4 Core E sum-rank

To analyze the relationships between *E*, *d*, and core-covariant association, we employed a multi-stage aggregation and ranking strategy. Initially, *E* was calculated for each core TSS and aggregated by summing the values for all TSSs sharing a unique *d*-rank. These totals were further aggregated by summing the *E* values of all TSSs falling within each specific *d*-percentile, consistent with the methodology used for core *d*-rank calculations. The resulting cumulative expression sums were sorted in descending order and assigned ranks, with the highest sum receiving the top rank.

A similar approach was applied to understand the association between *E* and the total number of covariants associated with a core TSS. *E* values were first summed for all core TSSs sharing a particular covariant count-rank. These values were then aggregated by summing all *E* totals falling within each specific covariant count-rank percentile. Following the same logic as mentioned above, these cumulative sums were sorted in descending order to assign final ranks, where the highest aggregate expression received the top rank.

#### 2.6.5 Mean E-rank

To examine the relationship between transcription cost coefficient and *d*, we calculated mean *E* by dividing the *E* of each core TSS by its total number of covariant partners giving rise to mean core-*E*. The *d* was processed by sorting the core TSSs in descending order, assigning ranks, and partitioning them into percentiles as mentioned above. For each *d*-percentile, we calculated the mean of these ratios across all core TSSs. These mean values were then sorted in descending order to assign final ranks. This mean *E*-rank is plotted on the Y-axis against the core *d*-rank on the X-axis to visualize how the average energy cost coefficient per covariant associates with genomic distance.

### 2.7 Calculation of Gini coefficient

The Gini coefficient was calculated and represented as a Lorenz curve to assess the distribution of transcriptional variables across the dataset. This analysis was applied to both the energy cost coefficient (*E)*, and the scaled CAGE scores. The Gini coefficient was derived using the following equation:

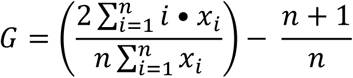

Where *n* represents the number of elements in the sample, *x_i_* denotes the value of each element, and *i* is the rank of the element when sorted in non-descending order. By plotting these values as a Lorenz curve, we visualized the cumulative distribution of expression and energy costs relative to the total population, providing a measure of inequality or concentration within the core-covariant networks.

### 2.8 Molecular function enrichment

To analyze the functional attributes of transcriptional connectivity, we performed molecular function enrichment analysis using GOrilla (Eden et al. 2009). For each core TSS, the associated genes were identified and subjected to ranked enrichment analysis. The core TSSs were ranked based on their mean core-*E* in both ascending order for performing enrichment and descending order, allowing for the identification of functional categories associated with varying levels of transcriptional energy cost. The resulting enrichment profiles are visualized as bubble plots.

### 2.9 Transcription factor binding site analyses of core TSSs

To analyze the enrichment of transcription factor binding sites (TFBS), we utilized the UniBind (Puig et al. 2021) database. For each Transcription Start Site (TSS), we extracted genomic coordinates in BED format, encompassing a region of 100 base pairs upstream and 100 base pairs downstream of the CAGE peak as curated by Satish et al. These 200-base pair windows were then used to identify and quantify the occupancy of transcription factor binding motifs within the selected regions of the core and covariant TSS populations.

#### 2.9.1 Differential enrichment of TFBS in core TSSs promoters over covariant TSSs

To identify transcription factor binding sites (TFBS) specifically enriched within the promoters of core TSSs relative to their covariant TSS populations, we utilized the UniBind Enrichment Analysis tool. The analysis was performed using the differential enrichment setting with the species parameter set to *Homo sapiens*.

This enrichment analysis was conducted independently for both robust and permissive TFBS collections across all model•data combinations. For each configuration, the top 10 differentially enriched TFBSs, as reported by the UniBind, were identified. This comparative approach allowed for the isolation of specific regulatory motifs that distinguish core TSS architecture from its associated covariant partners.

#### 2.9.2 Differential enrichment of TFBS in core TSSs promoters across different models (between N•N and C•C)

To identify regulatory motifs that are unique, we performed a comparative TFBS enrichment analysis using core TSSs that exhibited no overlap with covariant populations. This exclusive core TSS set comprised 725 sites for the non-cancer (N•N) configuration and 3,335 sites for the cancer (C•C) configuration. To account for the potential bias introduced by the higher population size in the cancer dataset, which might artificially increase the probability of motif discovery, we implemented a subsampling strategy. Specifically, we performed 10 random draws from the cancer core TSS pool, with each draw matched to the size of the non-cancer pool (n = 725).

Differential enrichment was then performed using UniBind, comparing the non-cancer exclusive core TSSs against each of the 10 randomly sampled cancer sets and vice-versa. This analysis was conducted independently for both permissive and robust TFBS collections, with the results subsequently merged. To ensure the stringency of our findings, we only retained TFBSs with a −𝑙𝑜𝑔_10_(𝑝) ≥ 2 that were consistently identified as enriched across all 10 iterations. For the final set of TFBSs satisfying these criteria, we calculated the mean occurrence count across all iterations to represent their enrichment magnitude.

#### 2.9.3 Differential enrichment of TFBS in core-TSSs promoters among same model, different data combinations

To investigate the enrichment of transcription factor binding sites (TFBSs) within core TSSs across different model•data configurations, we employed a two-stage strategy utilizing the UniBind extraction and enrichment tools. First, TFBS counts were extracted for both robust and permissive collections and subsequently merged. To ensure comparability across configurations, the raw counts for each TFBS were normalized by the total number of core TSSs identified in each specific model•data combination. For instance, counts for the N•C configuration were normalized by a factor of 0.59, while C•N counts were normalized by 0.69.

Enrichment was assigned to a specific model•data pool based on the positive difference in normalized counts between comparative pairs. For these assigned TFBSs, fold enrichment was calculated using a 𝑙𝑜𝑔_2_ transformation:

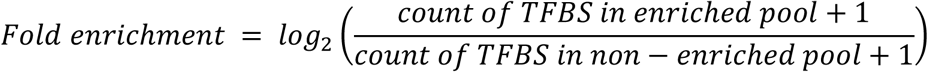

In the second stage, we utilized UniBind’s differential enrichment setting to determine the statistical confidence of these motifs. Differential enrichment was performed by comparing datasets processed through the same model architecture (e.g., N•N over N•C and vice-versa; C•C over C•N and vice-versa). These analyses were conducted independently for permissive and robust sets before merging the results. To maintain high stringency, we retained only those TFBSs with a maximum −𝑙𝑜𝑔_10_(𝑝) as 2. Finally, the previously calculated fold enrichment values were mapped to this filtered set of high-confidence transcription factor binding sites.

### 2.10 Targeted TSS-specific TBFS extraction and mean core-*E* comparisons

Transcription factor binding sites (TFBSs) identified as enriched in core TSSs across different data types using the same model architecture were filtered using a significance threshold of −𝑙𝑜𝑔_10_(𝑝) > 3. For these high-confidence motifs, specific TFBS instances were extracted using FIMO, utilizing the latest JASPAR collection (Ovek Baydar et al. 2026). The extraction was performed using the previously established BED region strategy, encompassing 100 base pairs upstream and downstream of the CAGE peak.

For each core TSS, we identified its associated TFBSs and calculated its mean core-*E*, defined as the ratio of core-*E* to its total covariant count and logged. This calculation was applied to all TFBSs identified across both consistent and cross-conditional model•data combinations (N•N, N•C, C•C, and C•N). Finally, the mean core-*E* values were plotted against each specific data pair to evaluate the relationship between transcriptional energy costs and TFBS occupancy.

### 2.11 Word letter diagram

To provide a comprehensive schematic of TFBS enrichment, we utilized a word-cloud visualization approach. In this representation, the scale of each transcription factor name corresponds to its statistical significance, defined by the −𝑙𝑜𝑔_10_(𝑝) as reported by UniBind. The TFBSs visualized in this manner were extracted following the methodology described previously and were filtered to include only those with a −𝑙𝑜𝑔_10_(𝑝) threshold greater than 3.

### 2.12 Statistical analyses and data visualization

Computational analyses were performed in Python (v3.13.2), utilizing a suite of scientific libraries for data processing and statistical evaluation. Data manipulation was primarily handled using NumPy, Pandas and Polars, while genomic interval operations were conducted via pybedtools and Biopython. Statistical significance was determined using SciPy and statsmodels, employing tests such as the Mann-Whitney U, Wilcoxon signed-rank, and Student’s t-tests, with p-values adjusted for multiple comparisons where applicable. Correlation analysis was performed using Pearson and Spearman coefficients.

Data visualizations were generated using Matplotlib, Seaborn, and Plotly to create high-dimensional plots. Specialized plots, such as Venn diagrams and word clouds, were produced using matplotlib-venn and the WordCloud library, respectively. All final figures and schematic diagrams were refined and composed using Inkscape.

### 2.13 Code availability

All computational analysis codes were written in Python (v3.13.2) and performed using Jupyter Notebooks. The analytical workflow is available on GitHub [access limited].

## 3 Results

### 3.1 Denoising of deep data for pan-promoterome TSS pairs reveals hierarchical hubs of co-deployments

The CAGE data was analyzed for understanding the structures of cooperative TSS deployments. We built upon the recent report by Satish and coworkers (Satish et al. 2025, under review) where autonomous TSS deployments have been reported to be different between curated sets of Cancer and Non-cancer samples. Using the same curated sets of 1255 Non-cancer (*N*) and 476 Cancer (*C*) samples we extracted features for TSS pairs instead of TSSs. For feature extraction we utilized three types of parameters: CAGE scores, TSS-start codon distance profiles and Shannon entropies associated with TSS deployments. For all the TSS pairs these 15 different features were calculated by combining CAGE scores with the distance between TSS and the most upstream start codon (*d*) and covariances. These 15 features represented a holistic set of dimensions in which TSS co-deployments (represented by covariances) related with transcriptional activity (TCI), *d* or both (*E*).

Features were derived separately for *N* and *C* sample sets resulting in a pool of 15 features for ∼1.5×10^9^ and ∼1.3×10^9^ unique TSS pairs for *N* and *C* respectively (Figure 1, A-C). The positions of the TSSs in each pair were fixed based on their *d* such that for two hypothetical TSSs A and B, a TSS pair A•B would always have *d* of A greater than that of B (more details in the method).

**Figure 1.**
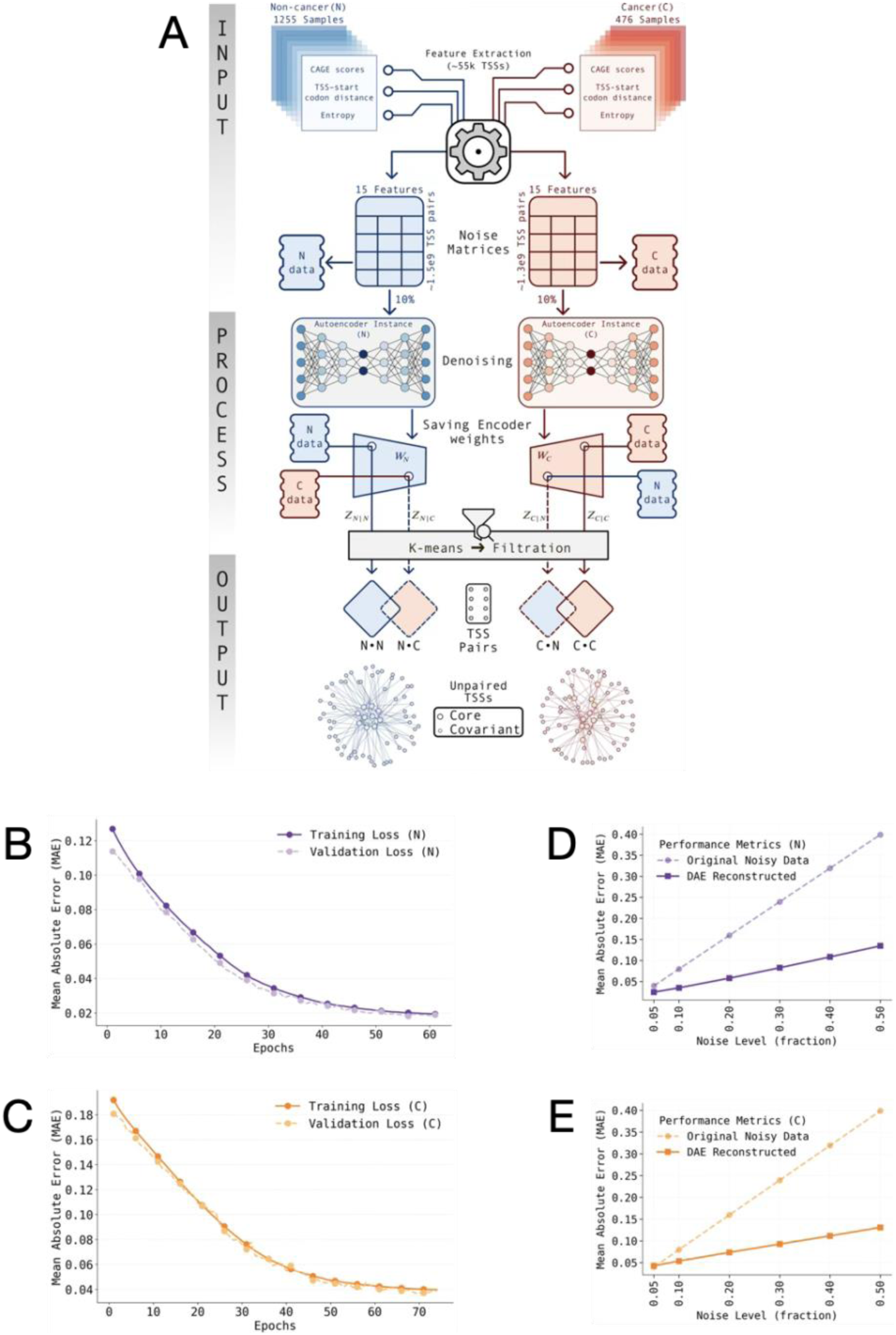
Analysis of “all TSSs by all TSSs” CAGE data matrices reveal TSS pairs with hierarchical co-deployments patterns with cancer and non-cancer distinctions. **A:** A schematic workflow describing the pipeline used for identification of core-covariant TSS pairs in CAGE data. A feature table comprising of 15 features (generated by combining TSS *d*, scaled CAGE scores and CAGE score-derived parameters Shanon entropy and inter-sample covariance) was derived for ∼1.4e9 – 1.5e9 unique TSS pairs separately for 1,255 non-cancer (*N*) and 476 cancer (*C*) samples. 10% of this data was used for training and validating a denoising autoencoder with a design that compressed the 15 features to 4 latent vectors (details in the methods section). The *N* and *C* denoising models thus generated were used to denoise the *N* and *C* CAGE data independently to latent vectors which were subjected to *k*-means clustering and the first clusters in each parsing with >100,000 TSS pairs were used for further analyses. The four exclusive sets of TSS pairs were thus obtained (stated as model•data these are N•N, N•C, C•C and C•N). Depending on the repeated occurrence of some TSSs as partners in the shortlisted pairs, unpaired TSSs were given the identities of core TSSs (smaller population, repetitive occurrence in pairs) or covariant TSSs (larger population, covarying with the core TSSs). **B and C:** The training and validation losses measured in terms of mean absolute errors (MAE) show that over the epochs in both training and validation losses converge by ∼60 epochs in *N* (**B**) and ∼70 epochs in *C* (**C**). **D and E:** The reconstruction fidelity of the *N* (**D**) and *C* (**E**) models was evaluated through a systematic perturbation analysis using additive Gaussian noise. Clean CAGE-derived input features were spiked with randomly generated Gaussian noise of increasing variance, represented as a noise fraction on the X-axis (ranging from 0.0 to 0.5). Denoising performance was quantified on the Y-axis by measuring the Mean Absolute Error (MAE) between the original ground-truth signal and the reconstructed output produced by the autoencoder. The results demonstrate significant model robustness, as both architectures maintained high reconstruction accuracy even at a 0.5 noise fraction. The divergence between the input error and the reconstruction error (delta MAE) confirms that the models have successfully learned the underlying data manifold, allowing them to actively filter stochastic noise rather than merely memorizing corrupted inputs.

These “15 featureśTSS pairs” matrices were subjected to dimensionality reduction using an autoencoder pipeline (details in methods). Random 10% draws of the two feature pools were used to generate autoencoder weights using a neural network of 2 hidden layers (12 and 8 dimensions) and a latent layer of 4 dimensions. The autoencoder weights for Non-cancers and Cancers were used to generate latent vectors for *N* as well as *C* data. The nomenclature of the derived latent vectors follows a “model•data” scheme henceforth (Figure 1A). The four sets of latent vectors were thus generated using different combinations of weights and data: *N* weights and *N* data (Z*_N_*_|*N*_), *N* weights and *C* data (Z*_N_*_|*C*_), *C* weights and *N* data (Z*_C_*_|*N*_) and *C* weights and *C* data (Z*_C_*_|*C*_). The four latent vector sets were subjected to unsupervised clustering (Figure 1A). The first cluster with 100,000 TSS pairs or more were selected for further analyses. Following the clustering, the shortlisted sets of TSS pairs for Z*_N_*_|*N*_, Z*_N_*_|*C*_, Z*_C_*_|*N*_ and Z*_C_*_|*C*_ were designated N•N, N•C C•N and C•C respectively (Figure 1A). Thus, the N•N and N•C were exclusive sets derived using *N* or *C* data respectively through *N* weights. Similarly, C•C and C•N were exclusive sets derived using *C* or *N* data respectively through *C* weights. The training and validation statistics showed that both autoencoder models *N* and *C*, although trained on independent data, exhibited highly similar Mean Absolute Error (MAE) loss patterns, plateauing after 60-70 epochs (Figure 1, B and C). We further assessed the models’ structural robustness using a Gaussian noise injection experiment. Given the extreme sparsity of CAGE signals, the MAE optimization deliberately drives the network to learn the stable baseline of the sequences rather than prioritizing high-fidelity reconstruction of extreme outlier peaks. Consequently, both models exhibited aggressive background denoising, significantly reducing the absolute error of the perturbed inputs. This confirms that the autoencoders effectively discard stochastic noise in favor of compressing the underlying biological structure into a robust latent manifold (Figure 1, D and E).

Interestingly, application of *N* weights to *N* or *C* data both gave rise to sets of co-regulated TSS pairs with unexpectedly skewed TSS compositions. A larger number of TSSs frequently paired up as covariants with a smaller set of TSSs. The core-covariant TSS count asymmetries appeared consistently in clusters regardless of the applied *k*-means counts (*k* ranging 3 to 20). We regarded the smaller set of TSSs as the core of a network in which the core TSSs pleiotropically pair with a larger covariant TSS set (Figure 2, A and B). The core versus covariant count asymmetry was comparable in N•N (Figure 2A) and N•C (Figure 2B), indicating that the core-covariant design is a feature of *N* CAGE promoteromes. In contrast, application of *C* weights to *N* or *C* data both generated a much less skewed distribution of core-covariant counts suggesting that the core-covariant count asymmetry is a feature of *N* promoteromes generally disrupted in cancers (Figure 2, C and D). The representative clusters selected for further analyses are highlighted (Figure 2, A-D). Interestingly, obtaining such a cluster was resisted the most by *N* data through *C* weights. We inferred from these findings that the *N* weights fetch an *N*-like property of CAGE data parsed through them. This property is a core-covariant model of TSS co-deployments, discovered with comparable propensities in *N* or *C* CAGE data. The *C* weights on the other hand fetch a distinct *C*-like property of the *C* CAGE data. The N•N, N•C, C•C and C•N core-covariant pairs were spread out throughout the genome with no apparent preference for any chromosome or region (Figure 2, E-H). The genome-wide distribution of core-covariant pairs shows a pan-genome disruption of the core-covariant mechanism in the pairs discovered through *C* weights (Figure 2, G and H) as compared to the pairs discovered through *N* weights (Figure 2, E and F). The *d*, covariance and *E* of the N•N, N•C, C•C and C•N TSS pairs were extracted and used for further analyses.

**Figure 2.**
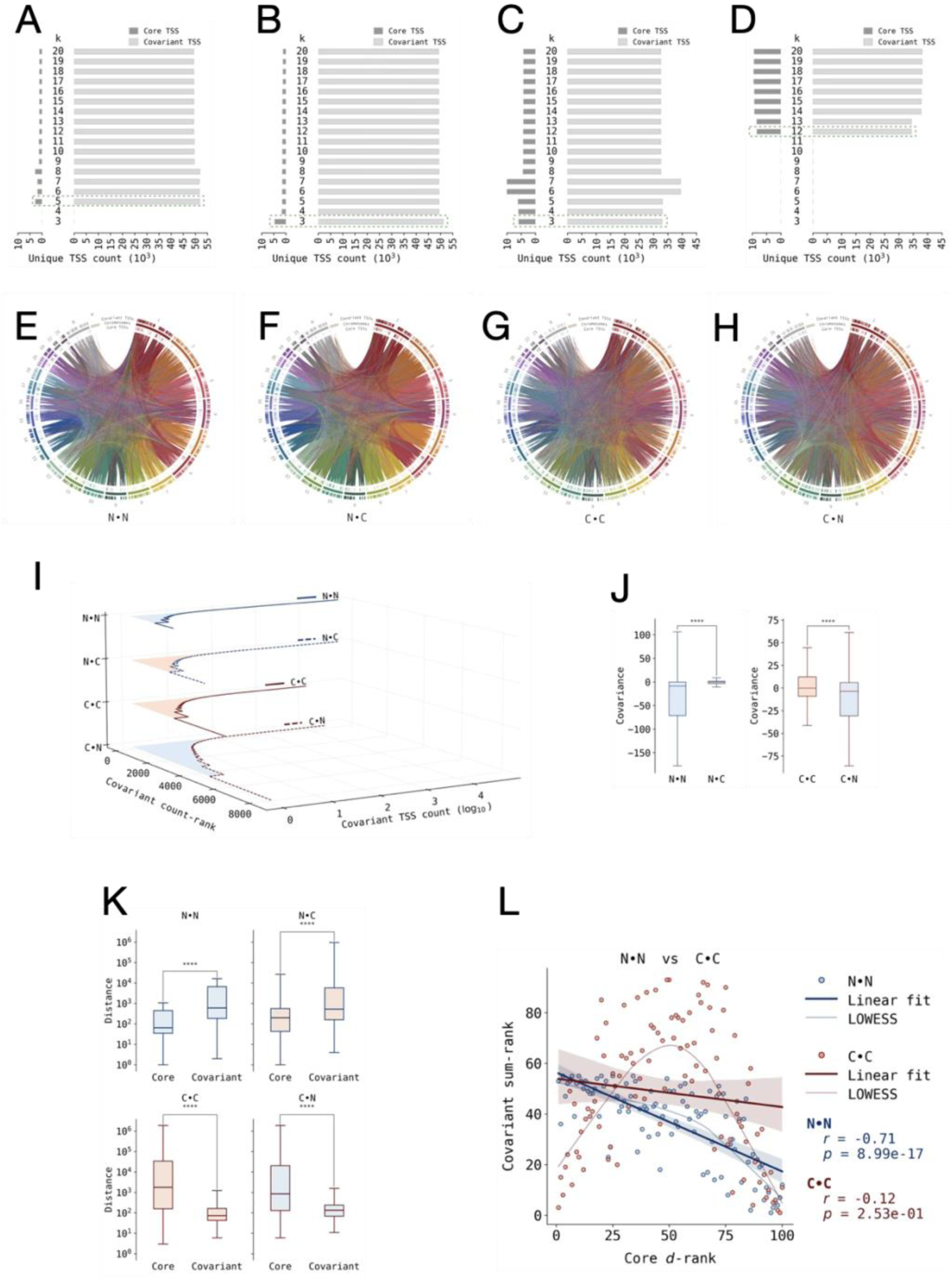
Pairs of co-deployed TSSs show a “one core TSS-to-many covariant TSSs” type of relationship genome-wide with widespread disruption in cancers. **A–D:** The core-covariant TSSs discovered as the largest non-noise cluster over multiple *k*-means iterations with increasing cluster counts are shown for N•N (**A**), N•C (**B**), C•C (**C**) and C•N (**D**). The horizontal axes are positive values in both directions from the split at zero and show the TSS counts of the core TSSs on the left (dark gray bars) and the covariant TSSs on the right (light gray bars). The intervening Y-axis shows the number of clusters in the *k*-means clustering iteration. The selection in green highlights the representative clusters selected for further analyses. The consistency of core-covariant count asymmetries is noticeable across the different *k*-means iterations. **E–H:** Circos plots showing the genome-wide landscapes of core-covariant pairs for N•N (**E**), N•C (**F**), C•C (**G**) and C•N (**H**). The peripheral circular axes show the chromosomes sandwiched between the covariant TSSs (outmost peripheral) and core TSSs (inner peripheral). The cores and covariant TSSs show a pan-genome distribution with no regional biases. The connectors between core and covariant TSSs denote pairs as identified in the selected k-mean clusters with color code being that of the covariant. The concentration of covariant color connectors at dense cores are visible in the model *N* (**E** and **F**) as against the *C* model (**G** and **H**). **I:** Core TSSs generated from the model *N* (N•N and N•C) have busier cores with higher covariant counts as compared to those generated by the *C* model (C•C and C•N). The distributions show core TSSs ranked by respective covariant TSS counts (descending) plotted against covariant counts. N•N and N•C show fewer ranks and higher covariant counts as compared to C•C and C•N in which lower covariant counts are shared between a larger number of core TSSs. **J:** Covariances between core-covariant pairs are governed by the CAGE data as well as the model. Despite the similarity in core-covariant counts, N•N and N•C showed extremely different covariance patterns with that in *N* CAGE data dominated by negative values and those in *C* CAGE data tending to zero. The core-covariant pairs identified by the *C* model showed a much broader distribution in C•C and C•N with positive covariances increasing significantly. **K:** In all TSS pairs identified through the *N* model the core TSSs were proximal to the respective start codons as compared to the covariants (distance on the Y-axis refers to the length of the genomic region between the CAGE peak center and the most upstream start codon for the corresponding gene as annotated in GENCODE). In contrast, for the TSS pairs identified through the *C* model, the core TSSs were distal as compared to the respective covariants (*p* < 1e-6, paired test). **L:** Core TSSs ranked by *d* (ascending rank order for descending *d*; X-axis) and covariant count (ascending rank order for descending count values; Y-axis) show that the N•N core TSSs exhibit an inverse correlation (*r_spearman_*=-0.71, *p*=8.99e-17) between *d* and covariant count (blue dataset) whereas the *d* and covariant counts are uncorrelated (*r_spearman_*=-0.12, *p*=2.53e-01) for C•C core TSSs (red dataset, insignificant linear regression fit). The C•C core TSSs exhibited a bimodal preference for high covariant counts at extremes of *d* ranks.

The relationships between a smaller set of core TSSs and a larger set of covariant partner TSSs could be used to understand the influence of non-autonomous mechanisms underlying TSS co-deployments. We focused on two primary features of the sets of TSS pairs: (i) covariances and (ii) covariant-core count ratios. Even though the covariances could be calculated only between pairs of TSSs, the repeated pairing between a core and multiple covariant TSSs suggested a one-to-many design of TSS co-deployments with multiple covariant TSSs corresponding to one core TSS. To understand and describe this pattern further, we calculated the covariant count for each core TSS. Even with a significantly less area under the curve (AUC), the N•N and N•C pairs accounted for co-regulation of >95% of all the 55,048 TSSs with as less as 2,722 and 4,577 TSSs at the core respectively (Figure 2I). In comparison, the C•C and C•N pairs had a larger AUC and yet accounted only for 35% and 64% of the entire TSS population although their core TSS populations were significantly larger at 5,759 and 8,298 TSSs respectively (Figure 2I). These results showed that a strong co-regulatory TSS core is characteristic of *N* and weakened in *C*.

The covariance profile of N•N TSS pairs showed the widest distribution (Figure 2J) with a preference for negative covariance. This could be explained by the tissue heterogeneity of the *N* sample set and the underlying cell type specific patterns of TSS deployments. The same *N* model applied to *C* showed a loss of covariance suggesting that the strong core-covariant design operates with an abundance of negative covariance in *N* promoteromes and no covariance preference in *C* promoteromes (Figure 2J). In contrast, C•C showed a range of covariances intermediary between those of N•N and C•C with a slight preference for positive covariances.

C•N showed a preference for negative covariances as well (Figure 2J). The most striking difference between the core and covariant TSSs was the difference in their *d* profiles. In N•N and N•C, the core TSSs had significantly lower *d* (Figure 2K). Conversely, in C•C and C•N the core TSSs had significantly higher *d* (Figure 2K). Next, we asked if the covariant TSSs preferred the core TSSs based on their *d*. Core TSSs were ranked by *d* (lower ranks for higher *d*) or the number of covariants (lower ranks for higher covariant counts). The core TSS *d*-ranks and covariant count ranks were significantly inversely correlated (*r_spearman_*=-0.71, *p=*8.99e-17) in N•N (Figure 2L). This correlation was lost in C•C due to a bimodal preference low as well as high d core TSSs for covariants (Figure 2L). Similar findings were obtained for N•C and C•N comparisons (Figure S1A) Thus, the *N* model identified core-covariant pairs where low *d* TSSs are preferred as cores by larger number of covariant TSSs and this preference for core TSSs was shifted towards higher d when the model *C* was used.

These analyses showed that the *N* model identifies TSS pairs with higher disparity in core TSS activities selectively. This selectivity of core TSSs is not a feature of the cancers and hence not observed if the model is trained on *C* data. We proposed that the nature of the core-covariant scheme in *N* represented a parsimonious design of TSS co-deployments due to the following features: A selectively active core, small core size with a large pan-promoterome approach, higher covariant count per core TSS. All these features confer parsimonious TSS deployments in *N* which are lost in *C*. To test this proposition, we studied these clusters further.

### 3.2 A parsimonious design with a minor set of pleiotropically co-deployed TSSs at the core is disrupted in cancer promoteromes

Parsimony of transcription starting at a given TSS can be considered at an individual TSS level (TSS autonomous) as well as at a systemic level. For the TSS-autonomous level, a parsimonious design would avoid futile transcription by preferring deployment of TSSs proximal to the start codons (low *d*) for any given gene (described by Satish et al). For the systemic level parsimony, we argued that (i) the core-covariant coregulatory structure would favor deployment of low *d* TSSs at the core and (ii) that the covariant TSSs would preferentially covary with the low *d* core TSSs. Thus, by preferring a low-cost core, the core-covariant scheme would attain parsimony. We analyzed the TSS clusters further to test if this proposition holds true and how this scheme differs between *N* and *C*.

When core TSS *E* were compared between all the four combinations N•N, N•C, C•C and C•N, we observed that *C* model or *C* data both associated with significantly higher *E* (Figure 3A). Since *E* combines TSS deployment with the futile transcription distance *d*, we concluded that the core TSSs operate through a low *E* scheme in *N* and a higher *E* scheme in *C*. These results (Figures 2L and 3A) showed that *d* as well as covariant profiles of core TSSs cooperate to minimize the resource costs of TSS co-deployments. Hence, as compared to the parsimonic scheme of core TSS deployment in *N*, the deviation in *C* is two ways: a preference for distal TSSs as well as an avoidance of low *d* TSSs. In order to test this more thoroughly, we compared the relationships between three different attributes of the core TSSS: *d*, *E* and covariant counts. Since these parameters have widely different ranges and non-uniform distributions, we used a ranked list approach (more details in methods). Core TSSs were ranked by their *d*, *E* or covariant counts and correlations between them were compared using linear regression. In N•N we observed that there was no clear correlation (*r_spearman_*=-0.09, *p*=3.61e-1) between *E* ranks and *d* ranks (Figure 3B). In contrast, a significant correlation (*r_spearman_*=0.40, *p*=3.08e-05) between *E* ranks and *d* ranks were observed in C•C showing that in C•C high *d* TSSs associate with high *E* (Figure 3B). A similar relationship between *d* ranks and *E* ranks were observed in N•C and C•N as well (Figure S1B), establishing that the association between high *d* and high *E* of core TSSs is not a CAGE data artifact rather a pattern fetched by *C* model and not by the *N* model. Similarly, correlation analyses between covariant count ranks and *E* ranks revealed that while N•N has no significant correlation (*r_spearman_*=0.12, *p*=2.19e-01) between these parameters, C•C has a significant preference (*r_spearman_*=-0.37, *p*=1.41e-04) for high *E* at core TSSs with low covariant counts (Figure 3C). The parsimony of the core-covariant design could be hence expected to manifest differently in C•C and N•N as core *E* per unit covariant (mean core-*E*). Correlations between mean core-*E* and *d* showed that the there is a significantly higher correlation between these two parameters in C•C than in N•N (Figure 3D). The core TSSs of model *C* operate with a higher *E* per unit covariant TSS from high *d* TSSs (Figure 3D). Similar observations were made by comparing core *d*, *E* and mean *E* of core TSSs for N•C and C•N suggesting that the observations made for C•C and N•N comparisons (Figure S1C and S1D) were truly due to the patterns identified by the models and not due to the data parsed through them.

**Figure 3.**
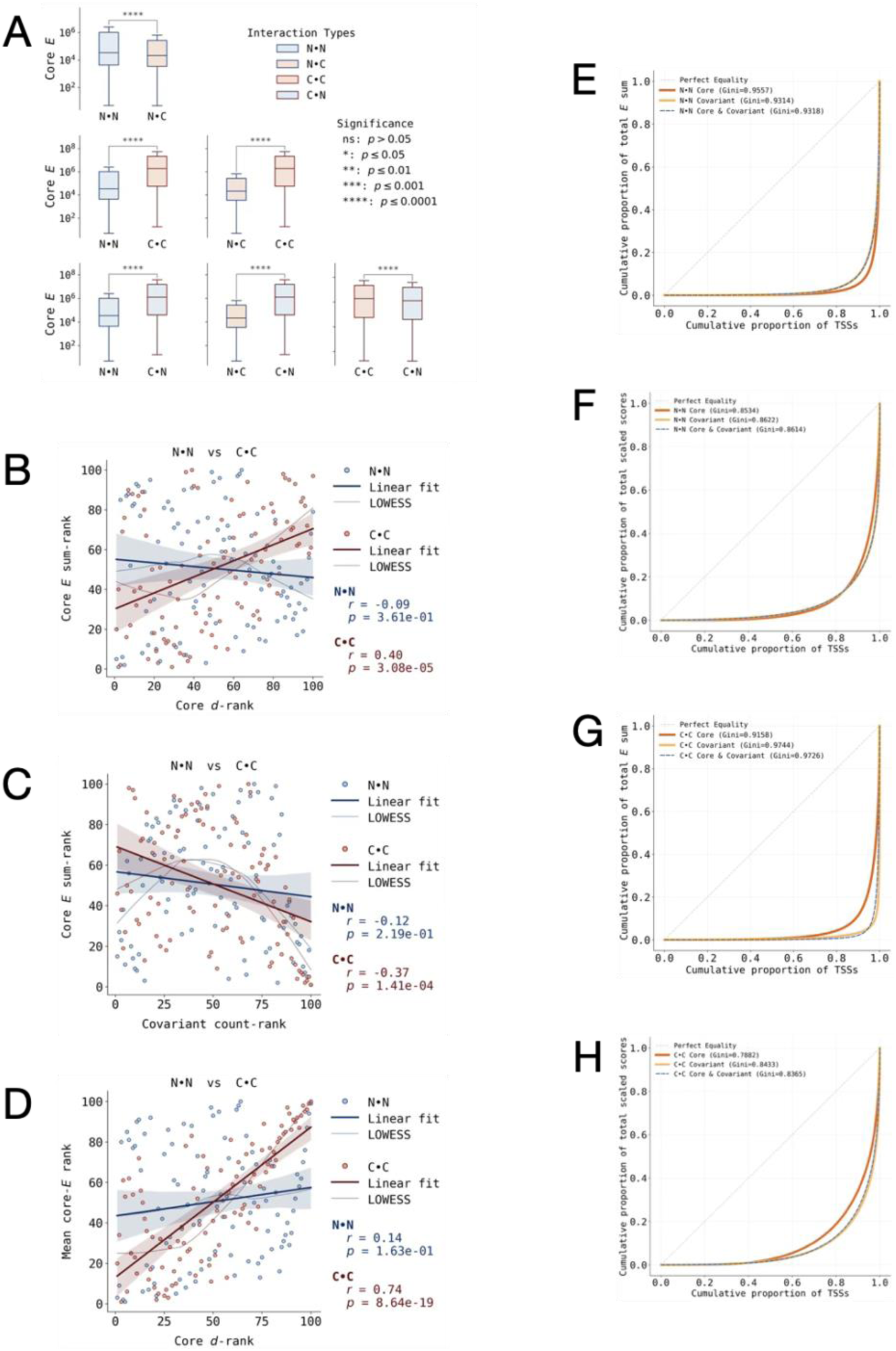
Non-cancer core TSSs exhibit parsimony through a preference for low *d* TSSs as covariant-dense cores which minimizes core *E* per unit covariant. **A**: *E* (the product of *d* and fractional CAGE score of a TSS) is significantly higher for core TSSs when any CAGE data (*N* or *C*) is parsed through the *C* model as compared to the *N* model. This *C* versus *N* model-dependence of high core *E* is reinforced by comparisons in which the models remain constant, but the change of the CAGE data increases *E* (N•C versus N•N and C•C versus C•N comparisons). **B**: The net core *E* ranks of the *N* model bear no correlation (*r_spearman_*=-0.09, *p*=3.61e-01) with core TSS *d* whereas the net core *E* for the *C* model is high at high *d* TSSs and low at low *d* TSSs with a strong correlation (*r_spearman_*=0.04, *p*=3.08e-05). **C**: Covariant count ranks and net core *E* ranks of the *N* model bear no significant correlation (*r_spearman_*=-0.12, *p*=2.19e-01) but the same in the *C* shows an inverse correlation (*r_spearman_*=-0.37, *p*=1.41e-04) such that *C* core TSSs with lower covariant counts have higher *E* and vice versa. **D**: The ranks of mean *E* per unit covariant TSS correlate with core *d* (*r_spearman_*=0.74, *p*=8.64e-19) for the *C* model such that as the core *d* increases, the mean core *E* also increases. This relationship is not significant (*r_spearman_*=0.14, *p*=1.63e-01) for the *N* model. **E-F:** An assessment of the cumulative distribution and inequality of *E* and scaled CAGE scores was performed for core-TSS populations in the N•N configuration. The core-TSSs exhibit a significant deviation from the line of equality (**E**) when evaluated by energy cost coefficient (*E*), suggesting a highly selective and concentrated distribution of transcriptional energy. In contrast, the distribution becomes more uniform when assessing CAGE scaled scores (**F**), indicating that while expression level is relatively distributed, the associated energy costs are governed by a subset of highly active sites. Calculated Gini coefficients for core-TSSs, covariant-TSSs, and the combined population are provided in the figure insets for quantitative comparison. **G-H:** In the cancer (C•C) state, core-TSSs demonstrate a distinct shift toward the line of equality when evaluated using the *E*. This trend suggests a reduction in selectivity and a more decentralized distribution of energy costs among core TSSs selected by the C•C model. Conversely, this distributional trend shifts when assessed using CAGE scaled scores, indicating that the C•C configuration exhibits altered regulatory selectivity. Calculated Gini coefficients for core TSSs, covariant TSSs, and the combined population are provided in the figure insets for quantitative comparison.

We next profiled the Gini distributions of *E* of the core and covariant TSSs to establish how TSS-selective or generic the nature of parsimonious co-deployments are. The Gini distributions of N•N, N•C, C•C and C•N TSS pairs were plotted for core and covariant TSSs separately as well as together. A comparison of the Gini distributions of *E* or the CAGE scores showed that the higher selectivity of core TSSs as compared to the covariants in the *N* model did not hold true in the absence of a *d* component. As compared to their respective covariant TSSs, the N•N core TSSs showed greater inequality (*Gini_E_*=0.96) in the distributions of *E* (Figure 3E) but not of CAGE scores (*Gini _Scaled scores_*=0.85) (Figure 3F) highlighting that the N•N core TSSs have selective *d*-dependent deployments. A nearly identical *E* and CAGE score Gini distributions were observed for core and covariant TSSs in N•C as well (Figure S2A and S2B). These results showed that selectively deployed core TSSs are a primary feature of denoised *N* CAGE data and although a similar set of *d*-selective deployment of core TSSs exists in *C* CAGE data, it is identifiable only by selectively denoising it through the *N* model. Gini distributions of *E* (Figure 3G) and CAGE scores (Figure 3H) in C•C showed a reduced selectivity of core TSSs (*Gini _E_*=0.92, *Gini _Scaled scores_*=0.7882). Similar observations could be made for C•N (Figure S2C and S2D). Thus *d*-selective *E*-conserving core TSS deployments turned out to be predominant features of *N* CAGE data and identifiable using *N* model readily in *N* or *C* CAGE data. A relatively less selective core TSS deployment turns out to be a more prominent feature of *C* CAGE data than that of *N*.

### 3.3 Cancer hallmark-supporting gene function enrichment of core TSSs is revealed by ranking using a parsimony parameter

The mean core *E* thus represented a parsimony parameter for the core-covariant co-deployments. These core-covariant schemes implied that the primary deployments of a minor set of core TSSs led to secondary regulated deployments of covariant TSSs akin to a signal amplification and diversification mechanism. Since the deployment of core TSSs also affects the associated gene function, we hypothesized that the genes represented by core TSSs must be hubs of specific functions.

Core TSSs were ranked by the mean core *E* in ascending or descending order, and corresponding genes were subjected to GO (BP) term enrichment analyses. Many GO terms were significantly enriched in core TSS lists ranked by mean core *E* in descending order showing that *E* per unit covariant is a functionally relevant parameter (Figure S3A) and that the core TSSs are not function-agnostic. Thus, the covariance between core and covariants appeared to have a structure in which the core TSS deployment is associated preferentially with certain gene functions. A large number of significantly enriched GO terms (FDR-corrected *p* value threshold 0.001) were exclusively enriched in N•N and C•N cores as compared to N•C and C•C respectively (Figure S3B). Amongst the GO BP terms enriched in the core TSS sets, the prominent ones corresponded to morphogenesis and differentiation, signal transduction, biosynthesis and metabolism, protein synthesis and degradation, transport and cell proliferation. Notably, TSSs belonging to cell proliferation genes were highly enriched in N•C whereas biosynthesis and metabolism were highly enriched in C•C (Figure S3C). Overall, the core TSSs derived from the *C* model also showed a preference for apoptosis and cell death related genes. No significant functional enrichment was observed when the same gene lists were sorted ascending for mean core-*E*, indicating that a function attribution of the core TSSs is energy and resource intensive and although quantitatively different, qualitatively very similar between *N* and *C*. Since the molecular functions of the genes associated with the core TSSs would be relevant to their effectiveness as cores, we next performed ranked enrichment tests for GO MF terms. Strikingly, all core TSS sets showed a very strong enrichment for RNA-binding molecular function when ranked by mean core-*E* descending (Figure S4A). When sorted ascending however, the most energy efficient core TSSs (with minimum mean core-*E*) were enriched in DNA-binding transcription factor activities with regulatory effects on RNA Polymerase II activities (Figure 4A). These were restricted only to N•N and C•C. The *C* model on the other hand showed exclusively albeit weaker enrichment of transmembrane transport of ions (Figure 4A). It was clear from these results that the molecular functions of DNA-binding proteins, especially transcription factors with RNA Polymerase II regulatory functions, are tightly linked to parsimonious functions of N•N core TSSs. We inferred that the hierarchical relationships between core and covariant TSSs operate through specific transcription factor binding sites.

**Figure 4.**
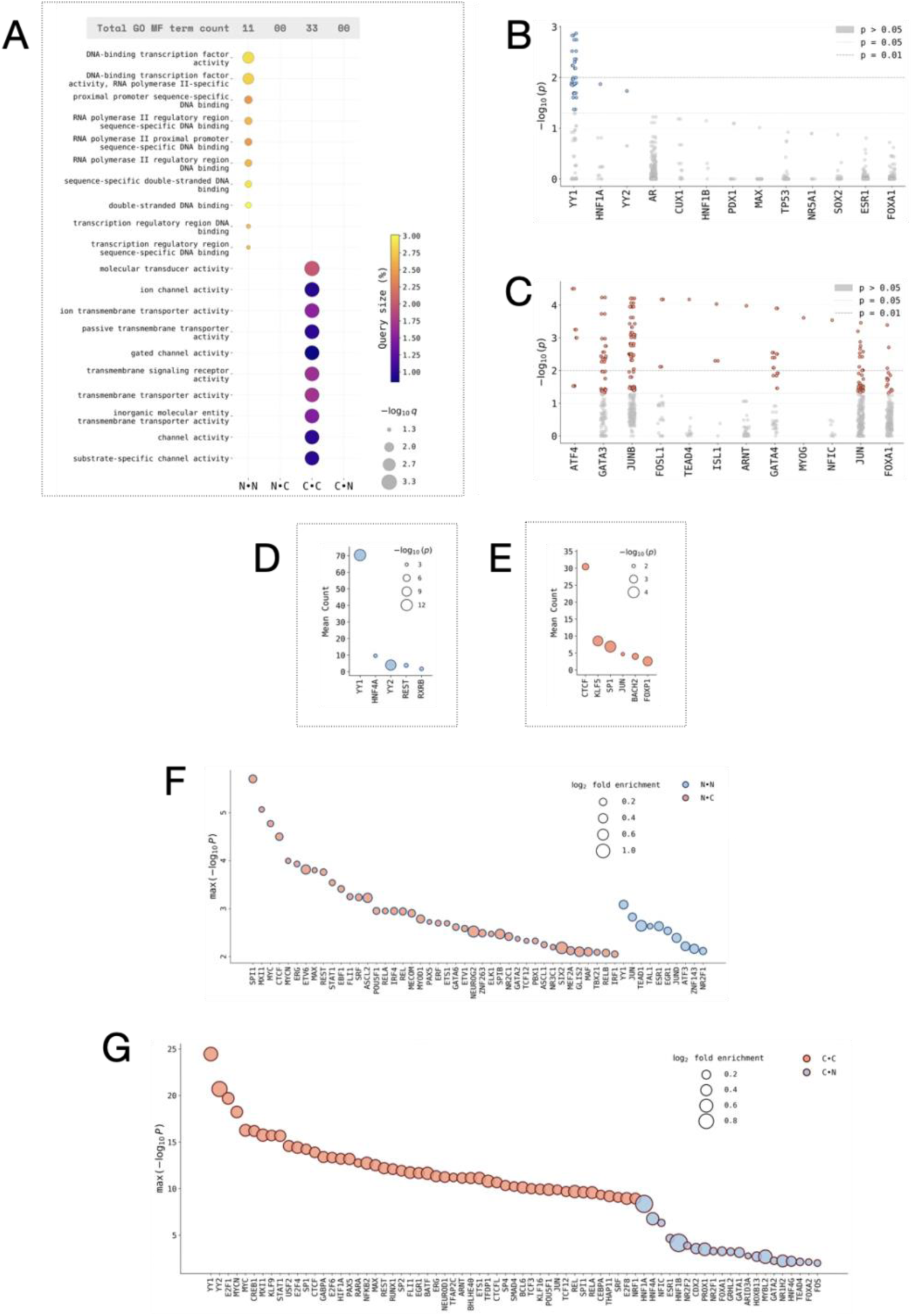
Core TSS genes ranked by mean core-*E* exhibit significant biological process and molecular function enrichment aligned with some cancer hallmarks. **A:** GO Molecular function (MF) terms corresponding to gene expression regulation were enriched in N•N when the core TSSs were ranked ascending by mean core-*E*. In contrast a similar GO MF term enrichment test for C•C showed enrichment only of terms corresponding to extranuclear and plasma membrane molecular functions supporting the possibility that transcription factor binding function associates with parsimony of core TSSs of the *N* model. **B and C:** Core TSSs were differentially enriched in specific TFBSs as compared to the covariants. The N•N core TSSs were enriched in three TFBSs, most prominently that of YY1 (**B**). The C•C core TSSs were enriched in multiple TFBSs including cancer hallmark supporting factors such as JUN and FOS (**C**). The Y-axes show -log10p for each enrichment instance separately. **D and E:** A direct comparison for differential enrichment of TFBSs in *N* core TSSs over *C* core TSSs (**D**) or *C* core TSSs over *N* core TSSs (**E**) shows that YY1 and CTCF are major enriched TFBSs in *N* core (**F**) and *C* core (**G**) TSSs respectively. **F and G:** Differential TFBS enrichments at N•C core TSSs compared to N•N core TSSs (red-filled circles) or N•N core TSSs compared to N•C core TSSs (blue-fille circles) with TF named mentioned below the X-axis (**F**). Notably the N•C cores show SPI1, MYC and CTCF as the topmost enriched TFBSs whereas a weaker enrichment in N•N cores are dominated by YY1 binding sites. A similar comparison between C•C (red-filled circles) and C•N (blue-filled circles) cores show enrichment of a much larger set of TFBSs dominated by YY1, YY2, E2F1 and MYC in C•C and multiple HNFs in C•N (**G**). The Y-axes show only the maximum -log_10_*p* for each TFBS out of all the enrichment instances, X-axes show discrete TF labels, color codes represent core TSS sets as indicated and datapoint circle area shows log_2_ fold enrichment in the indicated dataset (**F** and **G**).

The differences between core and covariant TSSs in terms of counts, *d* and *E* asymmetries suggested that the core TSSs function akin to a signaling hub where the transcriptional co-deployment is executed at target covariant TSSs. Since the core TSS identities varied widely between N•N versus N•C and C•C versus C•N, it was also obvious that the non-cancer to cancer differences of core TSS activities were less likely due to differential regulation of same core TSSs and more likely due to different inherent properties of core TSSs built into their sequences. For this possibility to hold true, it was necessitated that the core and covariant TSSs possess different DNA sequence properties such as regulatory protein binding sites. To test this hypothesis, we analyzed the TFBS profiles of cores and covariant TSSs in a narrow range of 100 base pair flanks from the CAGE peaks. Experimental data-based curated TFBS profiles were obtained using UniBind for core and covariant TSSs. The N•N and C•C core TSSs showed significant enrichments of different sets of TFBSs (Figure 4, B and C). Binding sites for the enhancer factor YY1 were almost exclusively enriched in N•N core TSSs as compared to the corresponding covariant TSSs (Figure 4B). The C•C core TSSs were enriched in TFBSs corresponding to multiple transcription factors including the activating factor ATF4 and activating protein components FOS and JUN. The long range trans-activating factors like YY1 were not observed in the C•C cores at all (Figure 4C). We investigated the differential TFBS enrichment as a regulatory feature of core TSSs.

### 3.4 The *cis*-acting enhancer-like TFBS profiles of core TSSs are dominated by YY1 and CTCF

In order to establish the consistency of TFBS differences in *N* and *C* core TSSs, we performed comparisons between core TSS sets in multiple different ways. We further performed detailed TFBS searches using UniBind (details in methods).

A differential TFBS enrichment between N•N core and C•C core TSSs showed that the YY1 enrichment is a marker for N•N core TSSs whereas the insulator factor CTCF is a marker of C•C core TSSs. Aligned with these findings, we observed that the core-covariant TSS covariances showed a distance dependence (Figure 4, D and E). To further understand this, we obtained TFBSs differentially enriched between N•N and N•C cores, and C•C and C•N cores (Figure 4, F and G). Reinforcing our other findings, we observed that N•C core TSSs were enriched in multiple TFBSs dominated by CTCF and MYC binding sites whereas N•N core TSSs were enriched in a much smaller set of TFBSs, most prominent of them being YY1 (Figure 4F). The C•C, unlike C•N, had core TSSs rich in a variety of TFBSs including YY1, CTCF, MYC and E2F1 suggesting a gross deregulation of core-covariant functionalities (Figure 4G). From these results we concluded that the identification of the core-covariant TSSs depended on the denoising pattern adopted by the *N* model differently from that adopted by the *C* model. These model-specific denoising led to identification of distinct core-covariant networks through *N* and *C* models based on *d* and *E* profiles. The CAGE data these models denoise offer TFBS-associated identities of core TSSs. The *N* data weighted through the *N* model represents the *N* core TSS setup marked by YY1 enhancer sites. The *C* data weighted through the *C* model represents the *C* core TSS setup marked by CTCF.

For all the core TSSs containing the differentially enriched TFBSs, we calculated the mean core-*E* independently from *N* as well as *C* CAGE data and compared them using Mann–Whitney unpaired tests. In N•C and N•N comparisons, the core TSSs containing binding sites for only two TFs, FLI1 and ELK1, showed higher mean core-*E* in cancers (Figure S5A) whereas those with binding sites for four TFs CTCF, NR2F1, EGR1 and SPI1 showed a lower mean core-*E* in cancers (Figure S5B). The mean core-*E* associated with these TFBSs showed no correlation between N•N and N•C suggesting a specificity of their deployments (Figure 5C). In a striking contrast, core TSSs with 51 TFBSs showed a significantly higher mean core-*E* in cancers (Figure S5C). Most interestingly, and as a reinforcement of the core TSS parsimony through DNA-binding proteins, none of the enriched TFBSs showed a reduction in mean core-*E* in cancers (not shown). These 51 TFBSs included transcription factors such as MYC, enhancer regulator factor YY1 and insulator factor CTCF. The loss of specificity of deployment of TSSs associated with these TFBSs was abundantly clear as their mean core E were highly correlated between C•C and C•N (Figure 5B).

**Figure 5.**
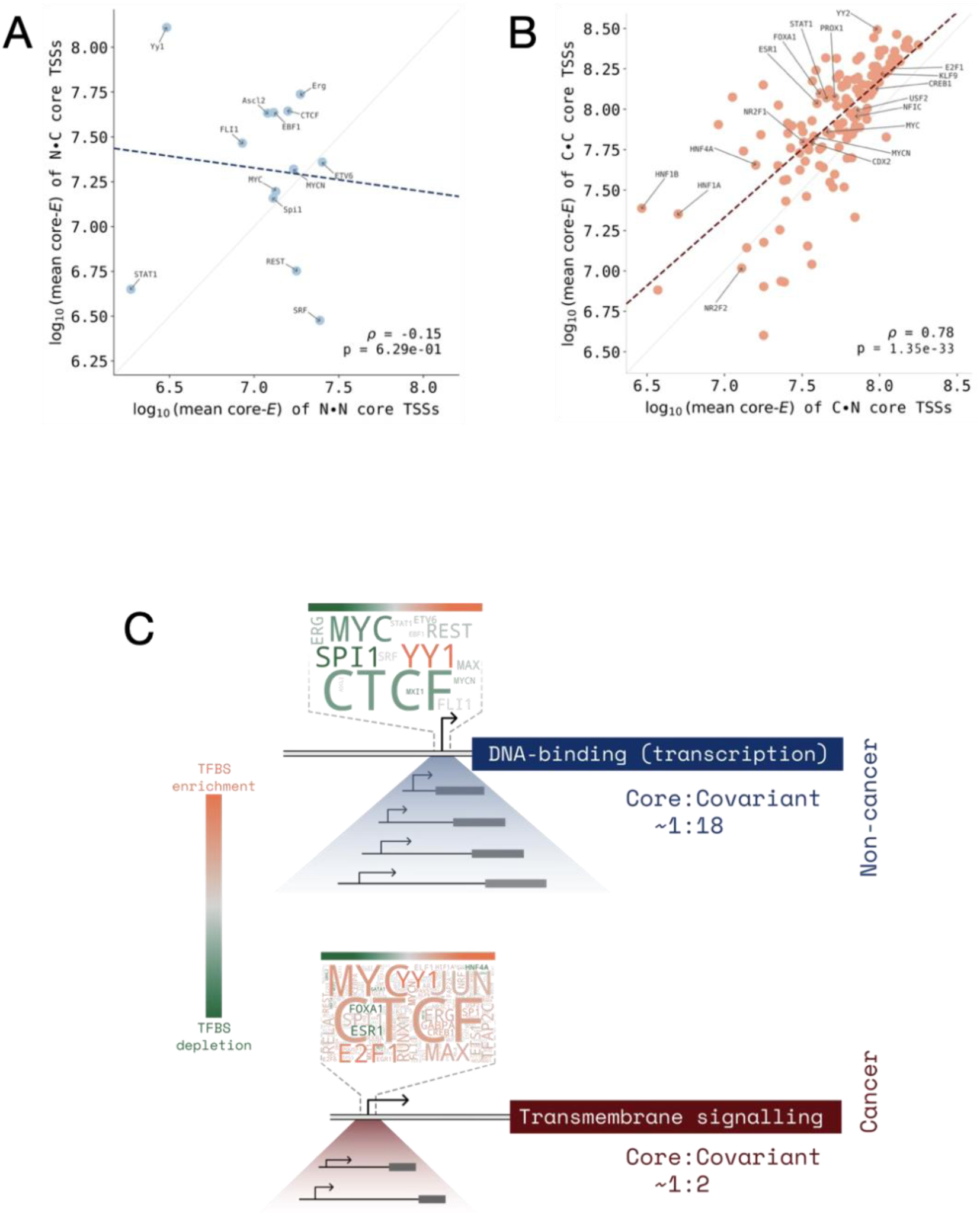
Mean core-*E* corresponding to differentially enriched TFBSs in *N*-core TSSs distinguish cancers and non-cancers whereas the *C*-core TSSs exhibit low TSS selectivity and cancer-non-cancer distinction. **A:** Mean core-*E* of N•N (X-axis) and N•C (Y-axis) TSSs show no correlation (*r*_spearman_=-0.15, *p*=6.29e-01) **B:** Mean core-*E* of C•N (X-axis) and C•C (Y-axis) TSSs show a strong correlation (*r*_spearman_=0.78, *p*=1.35e-33). The indicated values in A and B are log_10_ mean core-*E* and correlations are Spearman. Top 10 enriched TFBSs are named and identified with arrows. **C:** A schematic diagram summarizing the findings. Non-cancer core TSSs associated with genes involved in DNA-binding transcription factor activities are rich in YY1 and depleted in CTCF and MYC with a high covariant-core ratio in accordance with a parsimonic core-covariant regulatory network. The cancer core TSSs on the other hand are indiscriminately enriched in a large number of TFBSs, notably CTCF, YY1 and MYC. These cancer core TSSs have poor covariant-core ratio in accordance with a loss of selectivity and parsimony of TSS coregulation.

The recurrent enrichment of YY1 in *N* and CTCF in C core TSSs suggested that the enhancer regulation by YY1 and insulator function regulation by CTCF respectively affect the core TSS activities in *N* and *C* respectively. We tested if the recurrent occurrences of YY1 in *N* and CTCF in *C* indeed were indeed linked to the two different models. Covariances were reduced with increasing inter-TSS distances for all core-covariant pairs located in *cis* (limit of distance 2Mb) except for negative covariant pairs in C•C. As a control, such an inter-TSS distance dependence of core-covariant covariances did not exist for *trans* pairs (Figure S6A-B).

As summarized in figure 5C (Figure 5C), these findings combined with the gene function enrichment described earlier collectively showed that core TSSs exhibit covariance with their corresponding covariant TSSs through a mechanism that is (i) sensitive to biological functions such as differentiation, biosynthesis and cell proliferation at the core, and is executed through a high *E* directed towards an RNA-binding function and low *E* transcription factor activities. Overall, we concluded that the core-covariant TSS paradigm executes a functional covariance of TSS deployments in a gene function sensitive and hierarchical manner (Figure 5D).

## 4 Discussion

FANTOM CAGE data (Noguchi et al. 2017; Kawaji et al. 2017) are a deep resource for ascertaining independent TSS deployment frequencies in different tissues or samples (Noguchi et al. 2017; Kawaji et al. 2017). The CAGE scores represent the end-stage readout for each TSS independent of the other TSSs. The CAGE scores thus represent the TSS deployments in an “autonomous TSS deployment” framework (Satish et al. 2025, under review). However, TSSs share resources and cooperate as well as compete for them in the confines of nuclei. The results of such inter-TSS cooperation or competition are potentially a function of the genomic sequence properties of the TSSs as well as the epigenetic marks regulating them. Stochastic TSS deployment mechanisms under these regulatory constraints give rise to an overall TSS deployment landscape characterized by CAGE. The CAGE scores do not directly convey the patterns of TSS cooperation which lead to tissue type specific deployment patterns. In this study we have proposed and put to a rigorous test the following idea: CAGE scores can be used to extract biologically meaningful hidden TSS deployment patterns. By using deep learning-clustering techniques. We show that specific patterns of TSS co-deployments are embedded in CAGE scores. We also demonstrate that the TSS deployment patterns are tightly linked to TSS-start codon distance (*d*), relative TSS deployment rates, transcription factor binding sites and gene functions. We describe a hub and spoke-like model of core and covariant TSSs and show that they are disrupted in cancers. Our approach is dependent on a curated classification of CAGE samples into unambiguous non-cancer or cancer types as described elsewhere (Satish et al. 2025, under review).

Characterization of TSS co-deployments suffers from a lack of direct experimental approach. Even if CAGE scores are considered to be TSS-specific and TSS-autonomous metrics, the CAGE promoteromes potentially contain information about co-deployment patterns of multiple TSSs as a group. Such information is expected because TSSs do not exist and operate in isolation. TSSs colocalize in 3D to share transcriptional resources such as enzymes and transcription factors at enhancers and transcription factories (Di Giammartino, Polyzos, and Apostolou 2020; Gorkin, Leung, and Ren 2014; Wang, Cairns, and Yan 2019). In addition, resources such as nucleotides are competitively shared between TSSs even when they do not colocalize. While epigenetic marks and local chromatin environments confound any simplistic formulations of the TSS co-deployments, the CAGE scores can be utilized to fetch patterns of co-deployments. The TSS deployments, if random and mere statistical artifacts, would not exhibit functional and DNA sequence properties. Any approach based on known gene functions or DNA sequence properties are bound to be biased and would eliminate unknown co-deployment patterns a priori. Our approach is unbiased and based on a universal pair-wise deployment assessment of all TSSs equally. We have considered a null condition that all TSSs have a co-deployment element with all other TSSs which can be deciphered by patterns of CAGE scores for TSS pairs across multiple samples. We have utilized as features a combination of parameters that include TSS CAGE score covariances and entropy of TSS deployments. An additional parameter we have used to generate the features for analyses include the distance (*d*) between TSS and the most upstream start codon for the corresponding gene. It has been shown that the product of *d* and CAGE scores represent a transcriptional resource cost coefficient (*E*) for each TSS and is increased in cancers. The TSS deployment in cancers optimizes *d* in a gene function dependent manner with cancer hallmark supporting genes (biosynthesis and cell proliferation) preferring high deployments from low *d* proximal TSSs. The ∼55k TSSs were paired to form ∼1.5e9 unique TSS pairs and 15 features were derived for all these pairs. This approach ensured an unbiased characterization of TSS co-deployment patterns where the term co-deployment patterns were not just CAGE scores but also the co-variability in their deployments across the sample space. This approach however was noisy and data-intensive, and the denoising approach resulted in a reduction of TSS pairs by up to 3 orders of magnitude.

The TSS pairs obtained through the denoising algorithm are not mere statistical artefacts. Several observations establish that the TSS pairs identified through our denoising protocol represent a biologically meaningful and functionally relevant design. First, the protocol when applied to *N* or *C* datasets generates grossly different pairs of TSSs. This is established through comparisons between N•N and C•C. Since the TSS deployment differences between *C* and *N* could also contribute to identification of distinct TSS pairs using the *N* or *C* models, we parsed both *N* and *C* datasets through both the models. Thus, in a four-way comparison between N•N, N•C, C•C and C•N, we can delineate the effects of CAGE data on TSS pair identification from that of the model itself. Two primary properties of the TSS pairs stand out as not randomly expected from a noise matrix: (i) repetitive deployment of some TSSs with multiple others giving rise to a count asymmetry, and (ii) a strong *d* difference linked with the count differences. The essence of repetitive deployment of low *d* TSSs as partners of multiple covariant TSSs in *N* and its inversion in *C* shows that the denoising protocol identified genuine TSS pairs with biological relevance. It is striking that in the *k*-mean clustering step we continued to observe these count and *d* asymmetries across multiple clusters for each data-model combination. Interestingly, the count asymmetries implied a larger covariant per core TSS identifiable through the *N* model with >95% coverage of the entire promoterome against less then 50% coverage identified through the *C* model. These differences between the two models arose even when the training structures were identical. Also, these differences persisted even when the models were cross challenged by input data (*N* for *C*, and *C* for *N*). Such a design of TSS co-deployments suggest two things: Non-cancer cells maintain a design of core-covariant pairs with vast coverage of the promoterome, cancer cells suffer from a disruption of this design.

The core-covariant design reflects a signal transduction type paradigm in which the small set of core TSSs operate a selective diversification and amplification of transcriptional programs. This paradigm operates differently in cancer and non-cancers. The core-covariant paradigm is efficient in non-cancers and operates through low-*d* low-*E* TSSs. Parsimonious design of TSS deployment is achieved in non-cancer cells through the core-covariant mechanism. The covariant TSS pairs could colocalize topologically. Although TSS colocalizations are highly variable from sample to sample and generic representations of *C* and *N* are hard to expect as signals in publicly available datasets, we tested this possibility by trying out HiGlass (not shown). Beyond topological colocalizations of TSSs, we interpret these findings as co-deployment patterns imposed due to resource sharing from the common pool by all the TSSs. Resource sharing from a common nuclear pool is a possible mechanism of TSS deployment covariance.

Properties of the TSSs not revealed to the model or CAGE data in any way were found to be highly significantly different between the models as well as the CAGE data. Beyond *d* and *E*, these include TFBSs and *E* associated with TFBSs. First, for any model, TFBSs distinguish core TSSs from covariant TSSs in a CAGE data dependent manner. Thus, different TSSs with different TFBSs are deployed as core TSSs in *N* or *C*. When the models were compared, we observed that these are dominated by YY1 in *N* and CTCF in *C*. An obvious inference to be drawn from these findings are that the activities of YY1 (enhancer-promoter looping (Verheul et al. 2020)) and CTCF (insulator function and maintenance of large topological domains (Verheul et al. 2020; Lu et al. 2016)) are respectively key to core TSS identities in *N* and *C* respectively. The covariance in *C* is limited and tends to near zero. It means that insulator functions are disrupted in cancers. In *N* however, enhancers marked by YY1 exhibit large negative and significant positive covariances. *N* is a mix of heterogeneous tissue types with distinct tissue type enhancers in action, and this heterogeneity could expectedly give rise to YY1 as the major core TFBS. In contrast, in *C* the major TFBS being CTCF suggested that distal TSSs reshape the chromatin topology in cancers generically, act as core TSSs and affect covariant TSSs. Since YY1 and CTCF binding sites are known effectors of gene expression in long-range *cis* as well as *trans*, these findings lend support to the possibility that the core TSSs with YY1 and CTCF binding sites function as a regulatory core in the TSS deployment scheme. Our results also show that the *E* profiles of the core TSSs and their functional profiles support the core-covariant paradigm and its parsimonious nature.

Majority of the core TSSs in *N* cores maintained low *E* whereas only a minority accounted for the majority of *E* share. In *C* cores however, this selective *E* sharing to a minority of core TSSs was lost. The *E* cost of core TSSs per unit covariant was also high in *C*. This was due to two independent factors; high *d* of core TSSs and low covariant count per core TSS in *C*. Cooperatively, these two factors establish parsimony of the core-covariant co-deployment in *N*. Even when core TSSs were split according to the driver TFBSs, the high mean core *E* in *C* was established with only two TFBSs showing low *E* in *C*. Interestingly, the TFBS enrichment of core TSSs in *N* was limited and smaller (presumably due to tissue type heterogeneity in the *N* sample pool) as compared to the large number of TFBSs found enriched in *C*, including oncogenic factors such as MYC and JUN. The increased *E* in cancer for such a large set of TFBSs suggested a large-scale disruption of core-covariant mechanism of TSS co-deployments at the expense of parsimony.

A major identifying feature of the core TSSs, which is informative about the mechanisms of the core-covariant co-deployments is the functional category enrichment. In a remarkable reinforcement of the importance of mean core-*E*, we observed that the core TSSs exhibit biological process enrichment only when ranked descending by mean core-*E*. An ascending rank list showed poor to no biological process enrichment (not shown). It follows from these findings that YY1 and CTCF binding site associated TSSs of genes involved in functions such as biosynthesis, cell proliferation, signal transduction and protein transport account for the largest *E* share in a set of TSSs acting as core. An additional reinforcement of this mechanism comes from the fact that RNA binding and protein binding are the only two functional groups enriched in descending rank order of core TSSs. A combined synthesis of these results is that (i) the core TSSs selectively operate at a high *E* of individual core TSSs, (ii) the high *E* core TSSs affect covariant TSSs in a mechanism dependent on transcription factors of which the prominent ones are YY1 and CTCF, (iii) through RNA-binding and protein-protein interactions the core-covariant co-deployments are executed, (iv) the high *E* of only selected functional groups of TSSs ensures a systemic and overall low *E* in *N*, and (v) this mechanism is disrupted in *C* such that the parsimony of core TSS deployments is lost. Our findings position TSS co-deployments as an important mechanism and a useful parameter to understand transcriptional regulation and its dysregulation in cancers.

## 5 Author contributions

Conceptualization, RM, ALS and US; methodology, RM, ALS and US; software, RM; validation, RM, ALS and US; formal analysis, RM, US; investigation, RM, US; resources, RM and US; data curation, RM and ALS; writing—original draft preparation, RM, US; writing—review and editing, RM, ALS and US; visualization, RM; supervision, US; project administration, US; funding acquisition, US. All authors have read and agreed to the published version of the manuscript.

## 6 Conclusions

- TSS deployment patterns are mutually co-responsive resulting in patterns of co-deployments identifiable in FANTOM CAGE data.
- TSS co-deployment networks typically exhibit analogy to signal transduction networks in a minor core TSS population and a major covariant TSS population.
- Non-cancer samples maintain a parsimonious core-covariant co-deployment pattern with core TSS marked by TFBSs such as YY1 and enriched in transcription regulatory gene function.
- Cancer samples exhibit a gross disruption of the TSS co-deployment scheme marked by losses of core-covariant TSS networks, their parsimony, functional enrichment.
- Cancer hallmark supporting TFBSs mark the anomalous core TSSs in cancer CAGE data.

## 7 Acknowledgements

The authors acknowledge the service and support from the ParamAnanta Supercomputing facility at IIT Gandhinagar, multiple open source tools and software (as indicated in methods) and inputs from Praveen Kumar (IITGN).

## Supplementary figures

**Supplementary figure 1.**
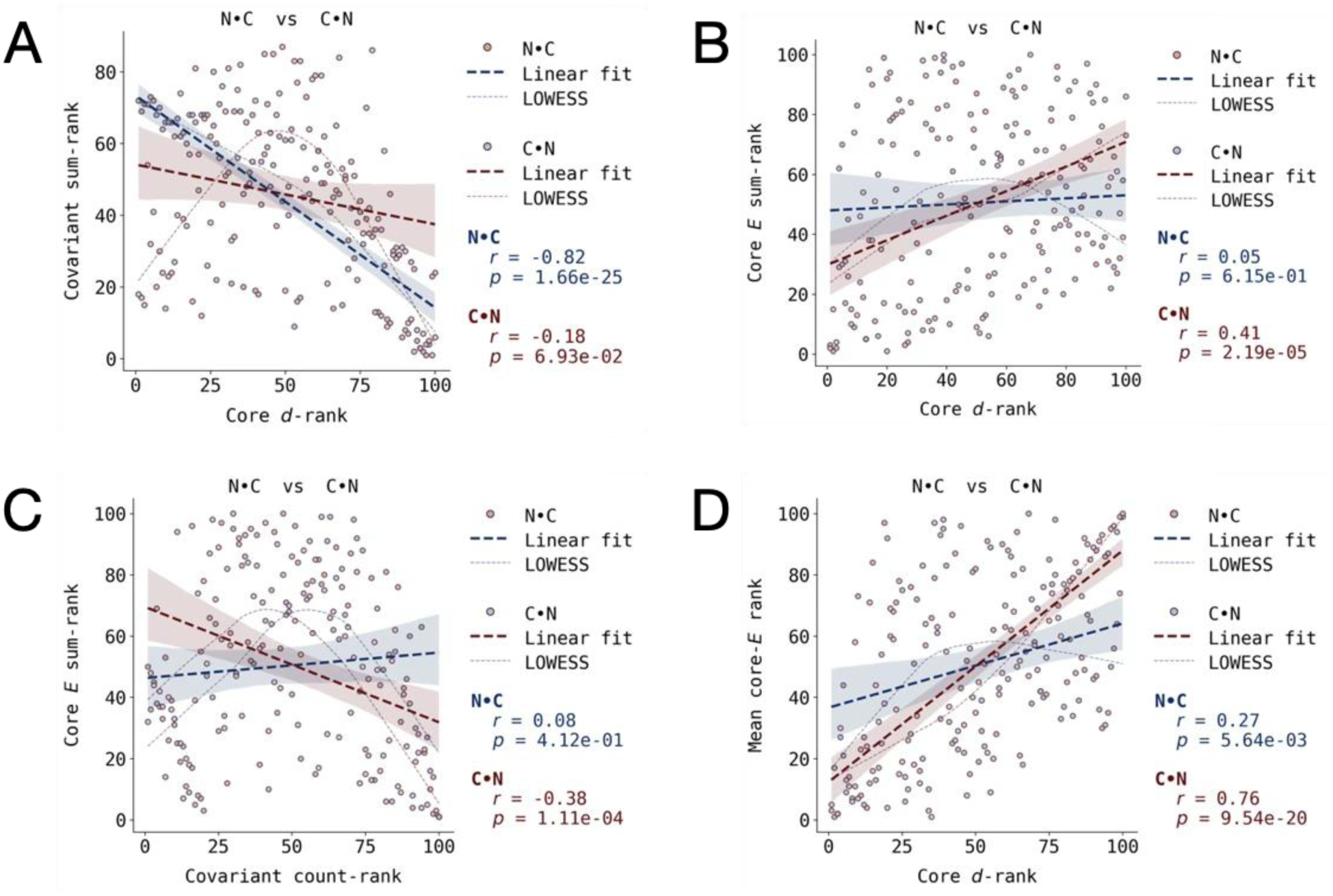
Core TSS-E and d can be utilized to demonstrate parsimony of core-covariant deployment. **A:** Core TSSs ranked by *d* (ascending rank order for descending *<u>d</u>*; X-axis) and covariant count (ascending rank order for descending count values; Y-axis) show that the N•C core TSSs exhibit an inverse correlation (*r_spearman_* = −0.82, *p* = 1.66 × 10^−25^) between *d* and covariant count whereas the *d* and covariant counts are uncorrelated (*r_spearman_* = −0.18, *p* = 2.53 × 10^−1^) for C•C core TSSs (red dataset, insignificant linear regression fit). The C•C core TSSs exhibited a bimodal preference for high covariant counts at extremes of *d* ranks. **B:** In N•C we observed that there was no clear correlation (*r_spearman_* = 0.05, *p* = 6.15 × 10^−1^) between *E* ranks and *d* ranks (B). **C:** In contrast, a significant correlation (*r_spearman_* = 0.41, *p* = 2.19 × 10^−05^) between *E* ranks and *d* ranks were observed in C•N. Correlation analyses between covariant count ranks and *E* ranks revealed that while N•C has no significant correlation (*r_spearman_* = 0.08, *p* = 4.12 × 10^−01^) between these parameters, C•N has a significant preference (*r_spearman_* = −0.38, *p* = 1.11 × 10^−4^) for high *E* at core TSSs with low covariant counts. **D:** Correlations between mean core-*E* and *d* showed that the there is a significantly correlation between these two parameters in N•C (*r_spearman_* = 0.27, *p* = 5.64 × 10^−03^) and in C•N (*r_spearman_* = 0.76, *p* = 9.54 × 10^−20^).

**Supplementary figure 2.**
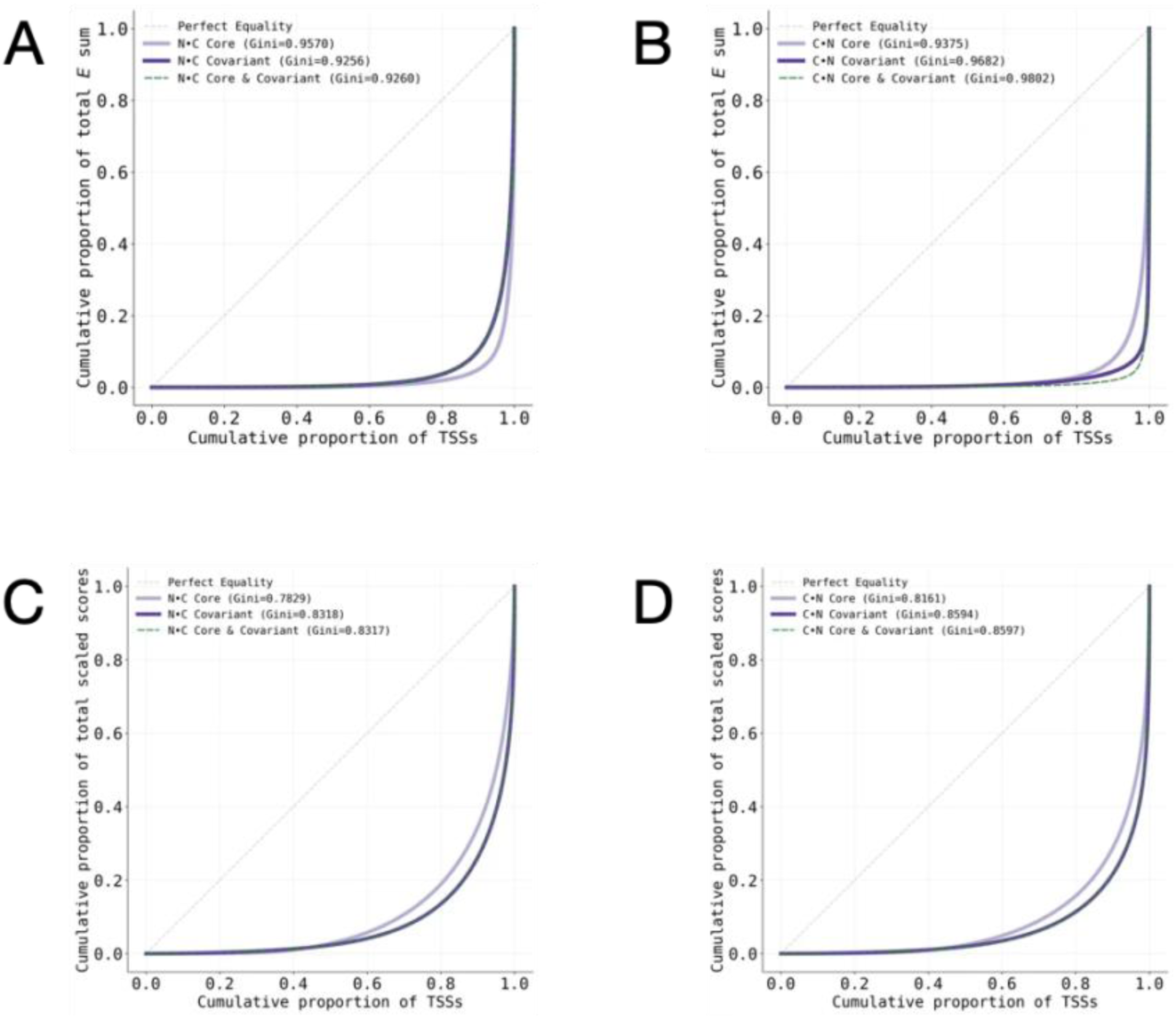
Assessment of selective *E* and scaled scores contribution by core and covariant TSSs. **A and B**: An assessment of selective *E* contribution by core and covariant TSSs in N•C and C•N using Gini distributions for the core TSSs and covariant TSSs. The core TSSs show a significant deviation towards inequality in N•C (**A**). In C•N however (**B**) the core TSSs exhibit an inverse shift towards equality suggesting that the core TSSs identified through the *N* model are selective whereas those identified through the *C* model are less selective in comparison. **C and D:** A similar assessment using the CAGE scaled scores for N•C and C•N does not show inverse deviations showing that *E*, and not just CAGE score, is key to the differences between the two models and the core TSSs identified through them. The Gini coefficients for core, covariants or both combined are mentioned in the inset.

**Supplementary figure 3.**
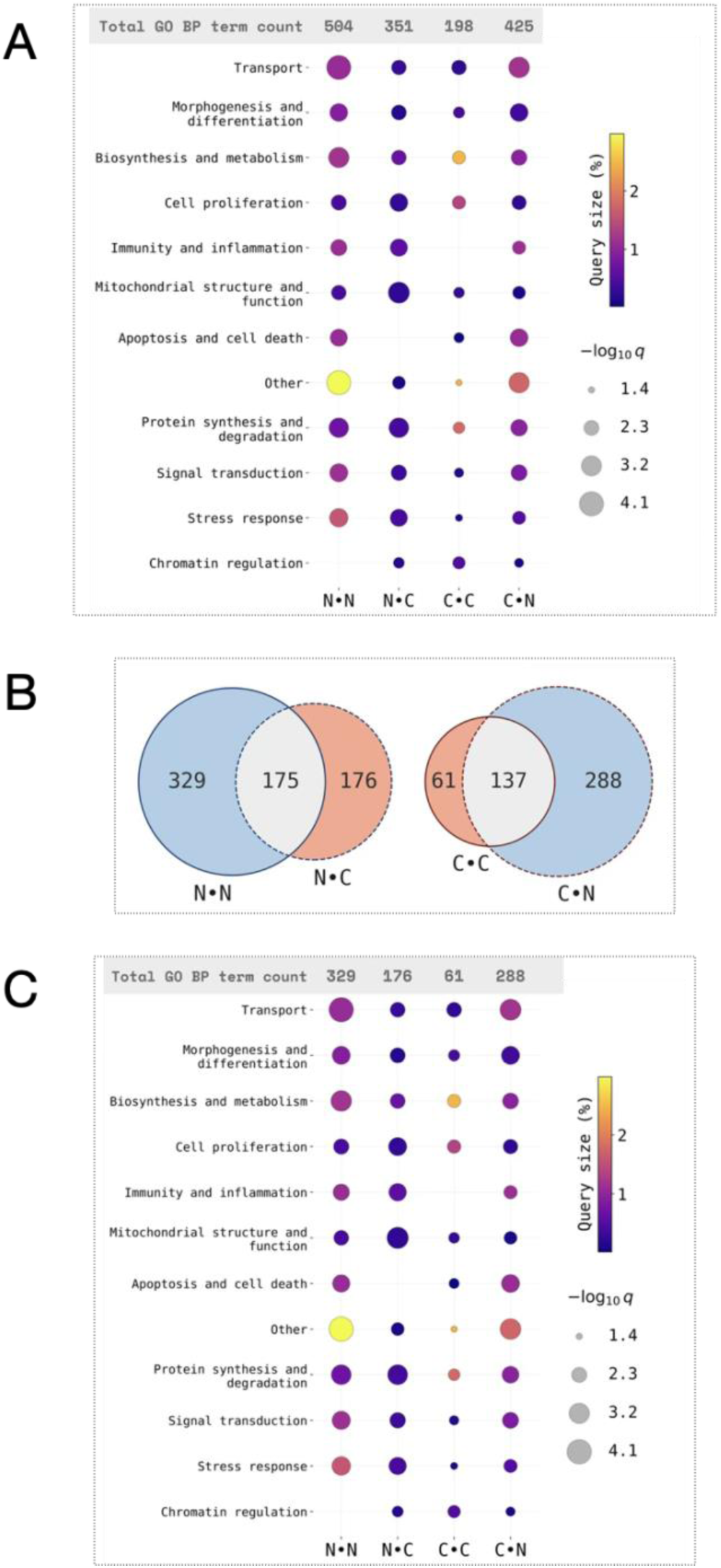
Genes associated with core TSSs when ranked by mean core-*E* show significant biological process enrichment. **A:** The core TSSs with high mean core-*E* (descending sort rank) are significantly enriched in GeneOntology Biological Process (GO BP) terms including cancer hallmark functions such as biosynthesis and cell proliferation. The actual GO BP terms have been condensed into representative blanket terms for representation. The actual GO BP term counts significantly enriched in each group is indicated at the top. **B and C**: Intersections between actual enriched terms in each of the four sets reveal that the *N* data generates core TSSs with more exclusive GO BP terms than the *C* data showing that the *N* model is more gene function-sensitive than the *C* model (**B**). The condensed GO BP terms occurring exclusively in N•N, N•C, C•C and C•N, with respective counts stated at the top, reinforces that the core TSSs are significantly associated with cancer hallmark-related GO BP terms (**C**). The GO BP term enrichments were observed only in high mean core-*E* (**A**–**C**) and not in low core-*E* (not shown).

**Supplementary figure 4.**
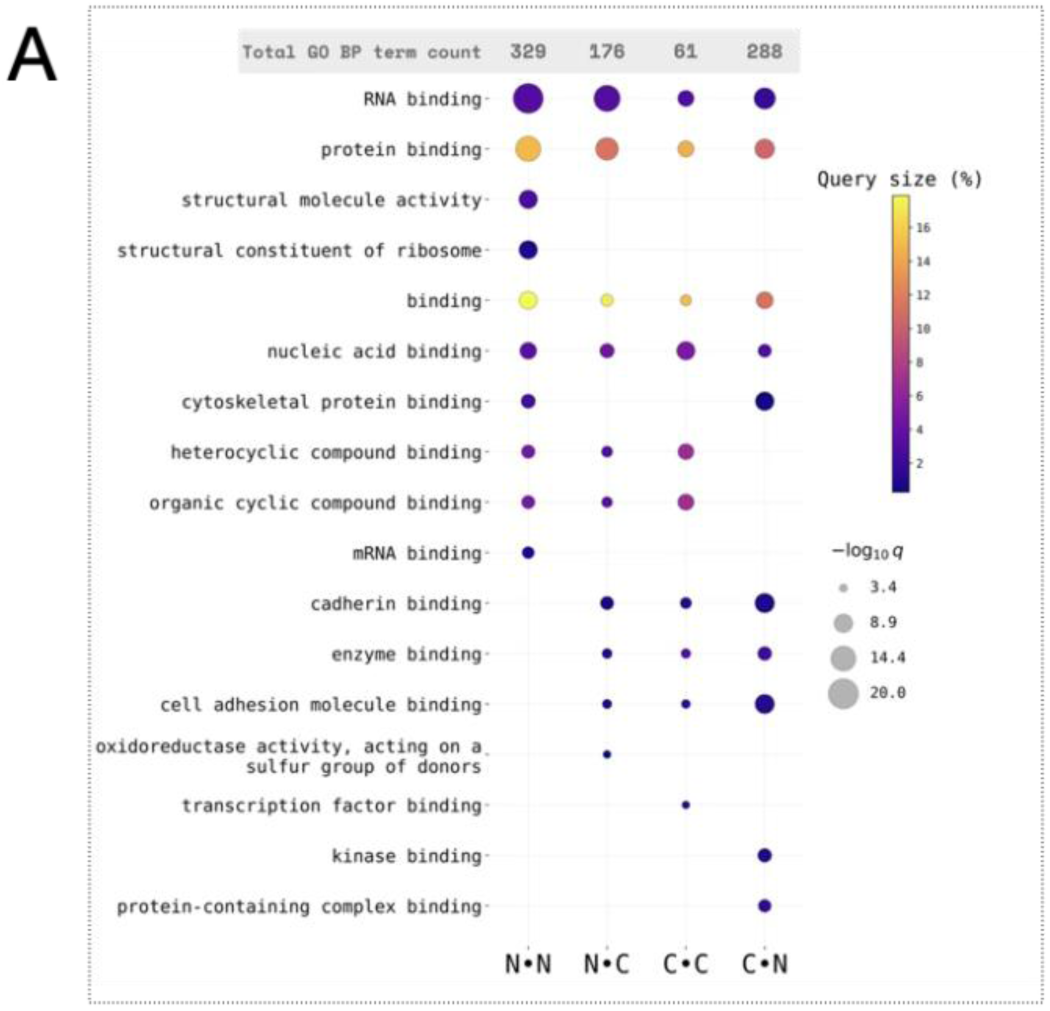
Genes associated with core TSSs when ranked by mean core-*E* show molecular function enrichment. **A:** GO Molecular function (GO MF) terms corresponding to gene expression regulation were enriched in high mean core-*E*. The high mean core-*E* TSSs of N•N, N•C, C•C and C•N were all enriched in multiple GO MF terms, especially RNA-binding.

**Supplementary figure 5.**
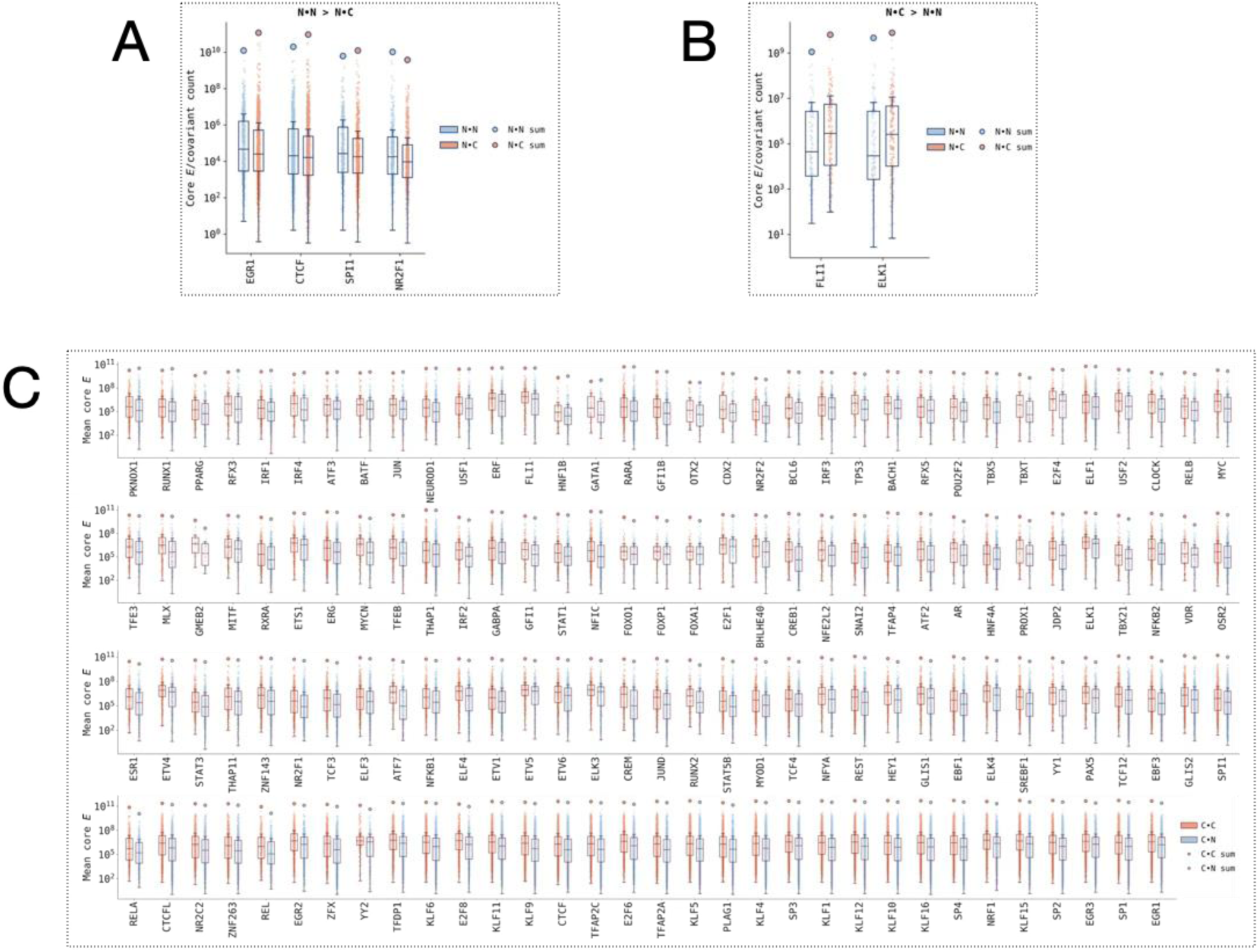
Non-cancer and cancer core TSSs are marked by distinct sets of TFBSs dominated by YY1 in non-cancers and CTCF in cancers. **A - C**: Mean core-*E* (core-*E* per unit covariant TSS) at *N*-core (**A** and **B**) or *C*-core (**C**) TSSs containing the indicated TFBS (x-axes) calculated from *N* CAGE (blue datapoints) or C CAGE (red datapoints) are shown as box-whiskers (the sum total *E* is plotted as a blob above the whiskers) for all the TFs with a significant difference between *N* CAGE and *C* CAGE data (unpaired comparisons). The mean core-*E* is significantly different between *N* CAGE and *C* CAGE only for *N* core TSSs containing six TFBSs (A and B), of which four (EGR1, CTCF, SPI1 and NR2F1) have a lower mean core-*E* in *C* CAGE whereas two (FLI1 and ELK1) have a higher mean core-*E* in *C*. In a striking contrast, all of the *C* core TSSs showed a mean core-*E* excess in *C* CAGE data and these could be mapped to be enriched in different TFBSs including CTCF, YY1, E2Fs and JUN amongst many other cancer hallmark supporting transcription factors (**C**).

**Supplementary figure 6.**
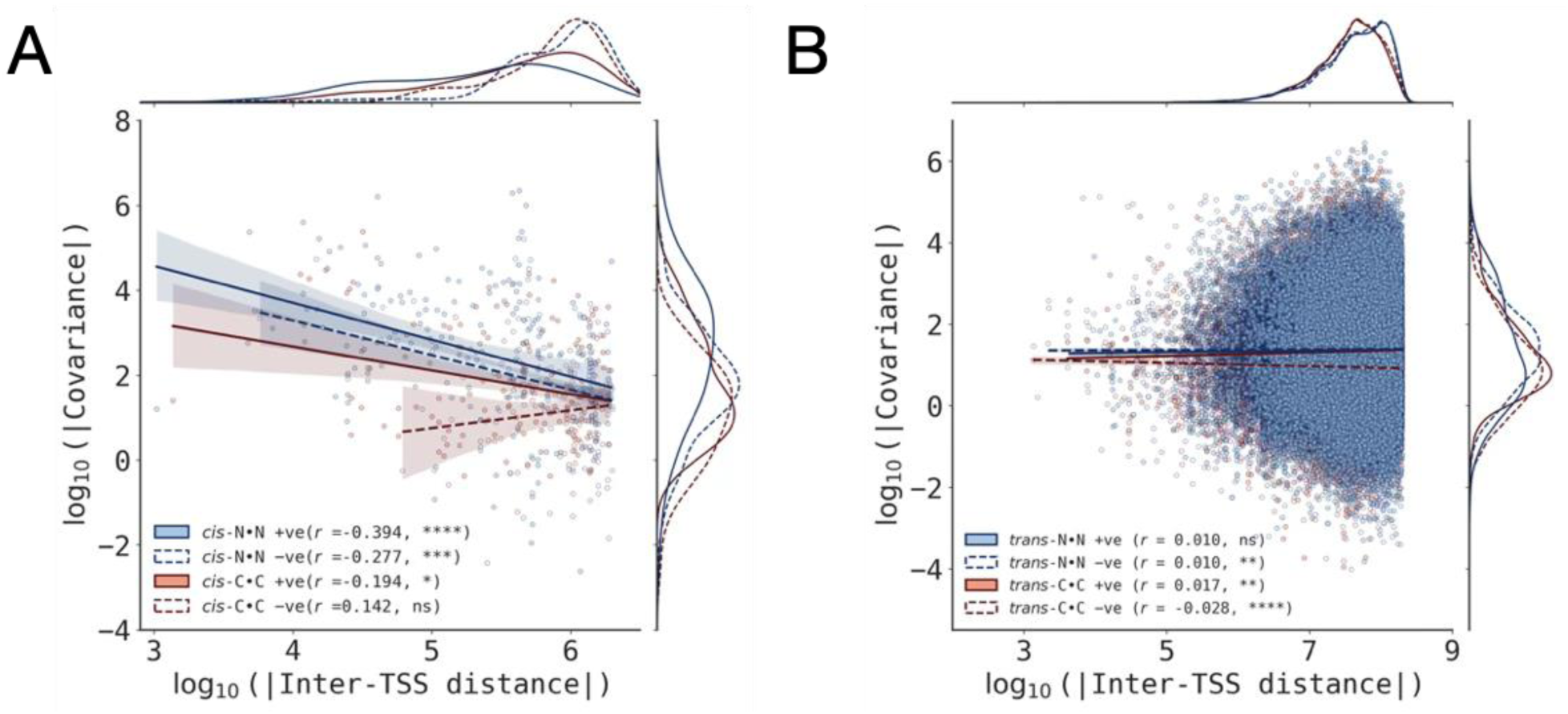
Covariance decreases with increase in inter-TSS distance for *cis*-TSS pairs. **A:** For N•N TSS pairs, both positive and negative covariance metrics decrease systematically as the inter-TSS distance increases. This relationship is supported by a statistically significant negative correlation between the absolute inter-TSS distance and covariance values. Conversely, C•C TSS pairs exhibit a distinct inter-TSS distance profile: they display a significant negative correlation exclusively with positive covariance, while showing an insignificant correlation for pairs characterized by negative covariance. **B:** Negative control visualization for trans-acting TSS configurations. Because trans-TSS pairs reside on different chromosomes or distinct genomic loci, a physical inter-TSS distance is biologically non-applicable; these pairs are mapped against a simulated distance metrics here strictly to serve as a baseline control to validate the *cis*-specific nature of the observed covariance decrease.

